# Tumor Cell Spatial Organization Directs EGFR/RAS/RAF Pathway Primary Therapy Resistance through YAP Signaling

**DOI:** 10.1101/2024.09.26.615226

**Authors:** Rachel Nakagawa, Andrew Beardsley, Sophia Durney, Mary-Kate Hayward, Vishvak Subramanyam, Nathaniel P. Meyer, Harrison Wismer, Hani Goodarzi, Valerie M Weaver, Daniel Van de Mark, Andrei Goga

## Abstract

Non-small cell lung cancers (NSCLC) harboring common mutations in EGFR and KRAS characteristically respond transiently to targeted therapies against those mutations, but invariably, tumors recur and progress. Resistance often emerges through mutations in the therapeutic target or activation of alternative signaling pathways. Mechanisms of acute tumor cell resistance to initial EGFR (EGFRi) or KRAS^G12C^ (G12Ci) pathway inhibition remain poorly understood. Our study reveals that acute response to EGFR/RAS/RAF-pathway inhibition is spatial and culture context specific. In vivo, EGFR mutant tumor xenografts shrink by > 90% following acute EGFRi therapy, and residual tumor cells are associated with dense stroma and have increased nuclear YAP. Interestingly, in vitro EGFRi induced cell cycle arrest in NSCLC cells grown in monolayer, while 3D spheroids preferentially die upon inhibitor treatment. We find differential YAP nuclear localization and activity, driven by the distinct culture conditions, as a common resistance mechanism for selective EGFR/KRAS/BRAF pathway therapies. Forced expression of the YAP^S127A^ mutant partially protects cells from EGFR-mediated cell death in spheroid culture. These studies identify YAP activation in monolayer culture as a non-genetic mechanism of acute EGFR/KRAS/BRAF therapy resistance, highlighting that monolayer vs spheroid cell culture systems can model distinct stages of patient cancer progression.

**Graphical Abstract:** 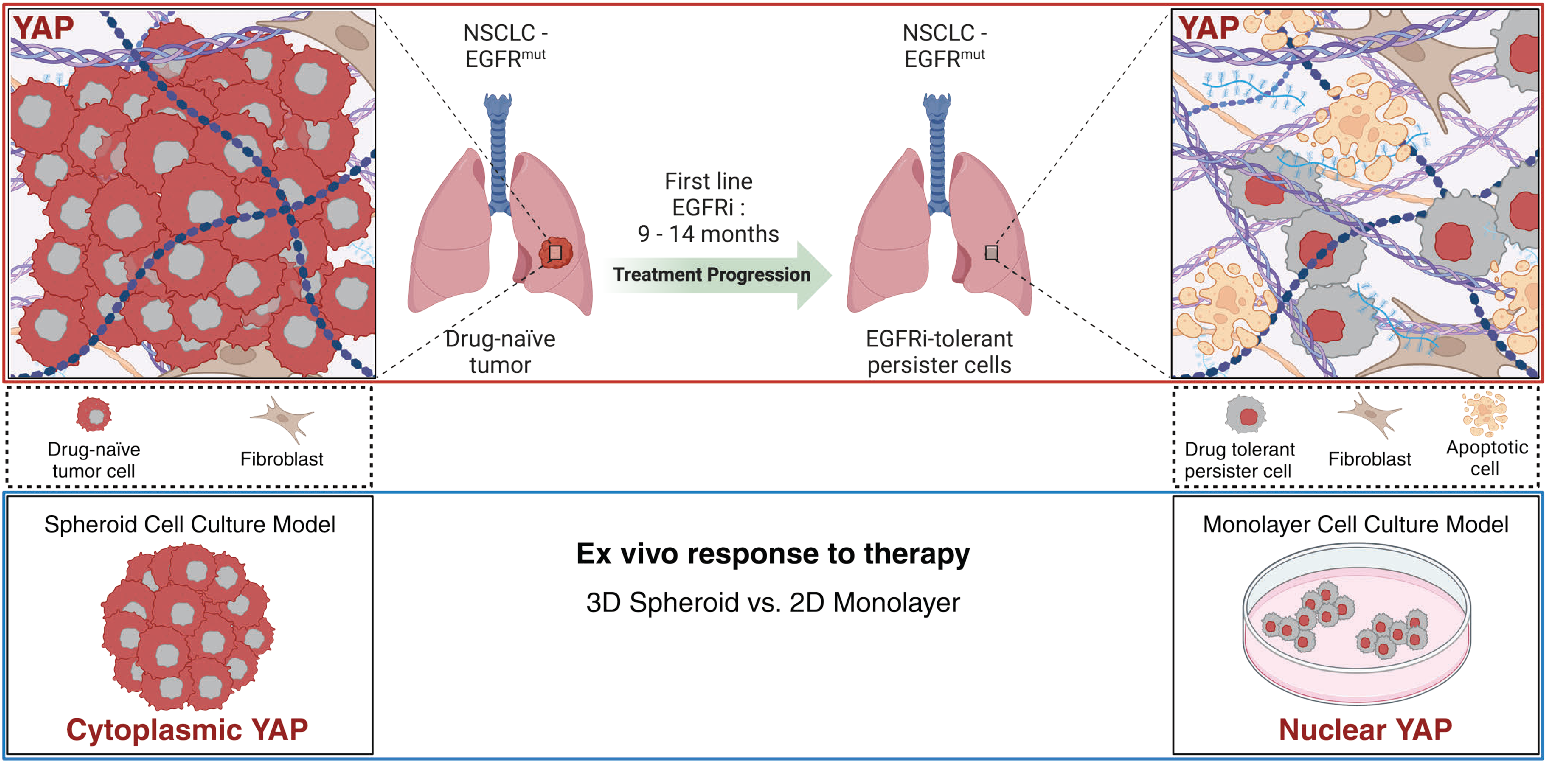

## Introduction

Non-small cell lung cancer (NSCLC) is the predominant lung cancer subtype in the United States, constituting approximately 85% of all diagnosed cases (1,2). Two frequently altered driver mutations in NSCLC are gene mutations in the proto-oncogene KRAS, which causes constitutive activation of the Ras pathway, or gene mutations in the epidermal growth factor receptor (EGFR). KRAS mutations are detected in approximately 30% of NSCLC patients. Second to KRAS mutations, approximately 19% of NSCLC patients will present with an *EGFR* mutation (1).

Small molecule EGFR inhibitors have transformed treatment for EGFR-mutated NSCLC (3), yet their initial tumor responses are often short-lived, with recurrence typically occurring within a year (4). Although newer generation EGFR inhibitors, including osimertinib, rociletinib, and olmutinib, target the T790M mutation, their efficacy remains transient (5,6), with no identified combinatorial therapies significantly prolonging EGFRi-mediated killing efficacy beyond monotherapy (7). Similarly, inhibitors targeting the KRAS^G12C^ mutation, such as ARS-1620, sotorasib, and adagrasib, bind to the mutant KRAS, resulting in temporary tumor regression in phase I/II trials, but with median progression-free survival averaging approximately 6 months (8– 12). Studies investigating resistance mechanisms of solid tumors to G12Ci uncovered KRAS^G12C^-mutant tumor cell populations use a multifaceted approach to bypass G12Ci-induced cell death (13– 16), with ongoing studies testing combinatorial therapies to improve G12Ci efficacy (17).

Though acquired genetic resistance mechanisms that allow cells to escape EGFRi or G12Ci are well studied, how tumor cells become acutely resistant to inhibitor-mediated cell death through non-genetic means remains poorly understood. Previous studies have implicated Yes-associated protein (YAP), a transcriptional co-activator that shuttles between the cytoplasm and nucleus in response to integrin signaling and biophysical stimuli (18–20), as a potential mediator of EGFRi and G12Ci resistance in NSCLC (21–24). Solid tumors develop and progress within a fibrotic stroma, characterized by extracellular matrix (ECM) remodeling and crosslinking that disrupts tensional homeostasis in epithelial cells and contributes to tumor progression and aggression (25– 28). Mechanical signals from a fibrotic collagenous ECM influence cellular responses, including YAP activation, which drives transcriptional programs promoting the proliferation and survival of tumor cells. The impact and dynamics of fibrotic tumor stroma on a subpopulation of cells known as drug-tolerant persister cells (DTPs) (29) that form following acute treatment and how cells responding to changes in tension may contribute to differential YAP activity in this cell population, however, remains to be explored.

Work identifying the role of YAP as a therapeutic resistance mechanism in vitro, however, may be confounded by the use of 2D polystyrene substrates used in traditional monolayer cell culture, as stiffer substrates have been shown to increase YAP nuclear translocation in mesenchymal stem cells and ovarian cancer cells (30,31). Polystyrene substrates are many orders of magnitude stiffer than tumor tissues and growth on plastic may alter cell shape and hyperactivate integrin-matrix adhesion and attendant downstream signaling (32,33). Constitutive integrin hyperactivation on plastic could modulate responses to targeted therapeutics, confounding interpretations of resistance phenotypes. To address this, multicellular tumor spheroids have been previously identified as an alternative cell culture method to better recapitulate in vivo physiology and chemotherapy response or resistance relative to monolayer culture (34–36). Here, we investigate YAP activity as a non-genetic resistance mechanism of EGFR/RAS/RAF pathway inhibition by comparing the acute effects of these drugs using in vivo xenografts in conjunction with parallel monolayer and spheroid NSCLC cell cultures.

## Methods

### Animal studies

Our study exclusively examined female mice because EGFR-mutant NSCLC is most associated with non-smoking Asian females (1); it is unknown whether the findings are similar between both sexes. To subcutaneously grow HCC827 tumors, the cells (1 × 10^6^) were subcutaneously injected into athymic nude mice (NCRNU sp/sp, Taconic Farms). Animals were enrolled simultaneously and treated with afatinib (Selleck Chemicals, S1011) dissolved in 0.5% methylcellulose/0.4% Tween 80 in water, at 20mg/kg, or vehicle alone. For evaluation of tumor remnant phenotypes, mice were treated daily for 5 days via oral gavage and euthanized on day 6. Tumors were harvested, fixed in 4% paraformaldehyde, and submitted for paraffin embedding (*n =* 4 vehicle; *n* = 4 afatinib).

### Antibodies and reagents

Drugs and inhibitors: afatinib (Selleck Chemicals), osimertinib (Selleck Chemicals), ARS-1620 (Selleck Chemicals), and vemurafenib (Selleck Chemicals) were all purchased solubilized in DMSO and used at indicated concentrations. Cisplatin (EMD Millipore) was dissolved in 0.9% NaCl w/w and used at indicated concentrations.

Antibodies for Western blots: β-actin (Santa Cruz #sc-47778), EGFR (CST #2232), pEGFR (CST #4407), YAP (CST #4912), YAP (CST #D8H1X), p-YAP (CST #13008), vinculin (CST #13901), Histone H3 (CST #9715). Restore western blot stripping buffer (Thermo Scientific). Ponceau S (Sigma).

Antibodies for IF: p-Histone H3 (CST #9706), pan-cytokeratin (sc-81714), YAP (CST #D8H1X), β-catenin (BD Biosciences #610154).

### Cell culture, cell lines, and virus production

HCC827, H1975, and H358 cell lines were obtained from the American Type Culture Collection (ATCC). HCC60 cells were a kind gift of John Gordon. RPMI-7951 cells were purchased from the UCSF Cell Culture facility. All cell lines were authenticated by STR and routinely tested negative for mycoplasma. Cell line culture mediums were as follows: 1) HCC827/H-1975 were cultured in high glucose DMEM supplemented with glutamine, pyruvate (Gibco), and 10% FBS (Gibco); 2) HCC60/H358 were routinely cultured in RPMI-1640 media (Gibco) and 10% FBS (Gibco); and 3) RPMI-7951 cells were culture in Eagles Minimum Essential Medium (EMEM) and 10% FBS (Gibco) at 37°C with 5% CO2. Spheroids were cultured using ultra-low attachment plates (Corning #7007). The pGAMA-YAP plasmid (Addgene plasmid #74942) and pGAMA-YAP-S127A plasmids were a gift from Diane Barber (58). Lentiviruses for FUCCI reporters and pGAMA-YAP-S127A were produced in 293T cells using standard polyethylenimine (Polysciences Inc.) transfection protocols.

### Picrosirius red staining and quantification

FFPE tissues were dewaxed in xylene and rehydrated in graded alcohols to deionized water. Tissues were counterstained with Weigert’s hematoxylin (Thermo Scientific, 88028 and 88029) for 10 min and then stained with 0.1% picrosirius red (Direct Red 80, Sigma-Aldrich, 365548 and picric acid solution, Sigma-Aldrich, P6744) for 1 hr. Polarized light images were acquired using an Olympus IX81 microscope fitted with an analyzer (U-ANT) and a polarizer (U-POT, Olympus) oriented parallel and orthogonal to each other. Images were quantified using a custom ImageJ macro to determine percentage area coverage per field of view using three to five fields of view per tissue. The ImageJ macro is available at https://github.com/northcottj/picrosirius-red.

### Trichrome blue staining and quantification

FFPE tissues were stained with Masson’s Trichrome (Thermo Scientific, 87019) according to the manufacturer’s instructions. Images were acquired using an Olympus IX81 microscope. Images were quantified using a color deconvolution ImageJ macro to determine percentage area coverage per field of view using three to five fields of view per tissue.

### Immunofluorescence staining and microscopy

Spheroids were consolidated from 96 microwell ultra-low attachment plates into a 15 mL conical tube, washed in phosphate-buffered saline with magnesium and calcium (PBS), and fixed in 4% paraformaldehyde at 4**°**C overnight. The spheroids were then washed three times in PBS and dehydrated using 30% sucrose in PBS overnight. The dehydrated spheroids were transferred to a Biopsy Tissue-Tek Cryomold (Sakura) and frozen on dry ice in Tissue Plus Optimal Cutting Temperature (OCT) Compound (Fisher HealthCare) and stored at −80 °C. Before sectioning, OCT frozen blocks were transferred to −20 °C. Using a standard protocol, 5 μm sections were cut from OCT blocks and affixed to Superfrost Plus microscopy slides (Fisher Scientific).

Tumor tissue was collected and fixed in 4% paraformaldehyde overnight. Using a standard protocol, tumor tissue was then dehydrated and paraffin-embedded, after which 5 μm sections were cut from paraffin blocks onto a warm water bath and picked up onto Superfrost plus slides. For immunofluorescence labeling, slides were dewaxed in xylene, rehydrated in graded ethanol (100%, 95%, 70%), and washed in deionized H20. Antigen retrieval was performed in 10mM citrate, plus 0.05% Tween 20 (EMD Millipore), pH 6.0 (Vector Labs), using a pressure cooker for 3 minutes.

For immunofluorescence of monolayer cultures, cells were grown on coverslips for 48 hours (fixed and permeabilized in 4% paraformaldehyde in PBS for 20 minutes at room temperature), followed by 3 PBS washes. Subsequently, monolayers and spheroids were blocked in IF buffer (1% bovine serum albumin and 2% fetal bovine serum in PBS) for 1 hour; tissue sections were blocked in 2.5% normal goat serum (Vector laboratories). Spheres, monolayer coverslips, or tissue sections were then incubated with primary antibodies diluted according to manufacturer recommendations overnight at 4°C, followed by 3 PBS washes, and incubated with Alexa Fluor-488 or -647 conjugated antibodies and counterstained with DAPI (Sigma).

Fluorescent images for tissue sections were acquired using a Zeiss ApoTome 3 equipped with an Axiocam 712 mono-megapixel digital camera using a Plan Apo λ 20x / 0.8 lens. Confocal microscopy was performed on an LSM 900 Airyscan 2 inverted microscope (Zeiss) using the EC Plan-Neofluar 10x/0.30 or PApo 20x/0.8 objectives.

### YAP immunofluorescence quantification

Quantification of YAP localization from tissues, coverslips, and mounted spheroid sections was performed using the StarDist segmentation extension v0.4.0 (cite 3 papers) generated in QuPath v0.4.3 using the training model *dsb2018_heavy_augment*.*pb provided by the StarDist authors*. In tissue sections, fluorescent cells were first segmented using the AF405-T4 channel collected from the Zeiss ApoTome 3. For monolayer and spheroid sections, fluorescent cells were segmented using the AF405-T4 channel (DAPI) imaged from the LSM 900 Airyscan 2 inverted microscope. Each cell detection provides fluorescent data from the nucleus and cytoplasm of identified cells. The “cell expansion” parameter in the StarDist extension was modified on a per sample basis to best fit the distance of the cell membrane past the nuclei as delineated by pan-cytokeratin, b-catenin, or EGFRdel19 membranous outlines. In tissue sections, after cells were segmented, cells were filtered by GFP+ for inclusion into analysis using the QuPath detection measurement manager to effectively exclude any non-tumor cells due to lack of pan-cytokeratin staining. Finally, intensity of YAP expression using the CY5 channel was determined, and YAP staining intensity in the nucleus was divided by YAP cytoplasmic staining intensity to generate a YAP positivity ratio. If the nuclear YAP / cytoplasmic YAP ratio was greater than 1.1 within a single cell, that cell was counted as having “nuclear YAP”. At least 5 sections across all samples were imaged and averaged across *n*=3 replicates.

### Differential YAP gene signature analysis

Microarray data from 83 matched pairs of lung adenocarcinomas and non-malignant adjacent tissue were retrieved from GSE75037 were retrieved using the GEOquery R package [https://academic.oup.com/bioinformatics/article/23/14/1846/190290]. The limma R package [https://academic.oup.com/nar/article/43/7/e47/2414268] was then used to perform differential expression between non-malignant and adenocarcinoma tissues. Genes were then ranked according to their t-statistic and the gsea enrichment analysis was performed using the fgsea R package [https://www.biorxiv.org/content/10.1101/060012v1] for the CORDENONSI_YAP_CONSERVED_SIGNATURE and YAP1_UP pathways from MSigDB. To perform Z-scored enrichment analyses, counts were first z-scored across samples. Expression scores for a gene signature were retrieved by averaging the z-scores of genes within the geneset, for each sample. These expression scores were then compared between conditions using a t-test.

### Western blotting

For cell lysis, Pierce RIPA buffer containing protease inhibitor (cOmplete Mini, EDRTA-free, Roche) and phosphatase inhibitor (PhosSTOP EASYpack, Roche). Monolayers were harvested by scraping cells directly into lysis buffer after 3 PBS washes and homogenized by passing 5x through a 27G needle. Protein extracts were measured using the DC protein assay (Bio-Rad) and normalized using the Pierce RIPA lysis buffer. Extracts were then boiled and resolved by electrophoresis in Bolt 4-12% Bis-Tris Plus gels and transferred to nitrocellulose membranes using the iBlot 2 Gel Transfer Device (Thermo). Quality control of the transfer was assessed by membrane staining in Ponceau S (Sigma-Aldrich). Membranes were then washed in TBS-T and blocked in 5% milk for 1hr at room temperature, probed with primary antibodies overnight at 4°C shaking, washed 3 times, probed with HRP-HRP-conjugated secondary antibodies for 1h at room temperature, washed 3 times, and finally incubated with Visualizer Western Blot Detection kit (Millipore) according to manufacturer protocol. Chemiluminescence was visualized using a ChemiDoc XRS+ imaging system with Image Lab Software (Bio-Rad).

### Viability and apoptosis assays

For monolayer relative ATP assessment, cell lines were seeded into 96 microwell TC-treated plates. 24 hours after seeding, cells were treated with assigned inhibitors or chemotherapeutics at the indicated dose. 72 hours following treatment, cells were incubated with Cell-Titer-Glo Viability Assay reagent (Promega) according to the manufacturer’s instructions. Luminescence was measured in a Tecan Safire^2^ plate reader. For spheroid relative ATP assessment, cell lines were seeded into a 96 ultra-low adhesion plate (Corning) for 96 hours, then treated with assigned inhibitor or chemotherapeutic at indicated dose for 72 hours. Following treatment, cells were incubated with Cell-Titer-Glo 3D Viability Assay reagent (Promega) according to manufacturer instructions. Luminescent signal was detected by a Tecan Safire 2 plate reader using Magellan analysis software. Experimental values were normalized to DMSO-control and GraphPad Prism 9 was used to transform concentration to log form and run non-linear regression (either four or three parameters) to generate best-fit values used for analysis. For apoptosis visualization, inhibitor- or DMSO-treated monolayers were treated with media containing 2X propidium-iodide (PI) and Hoechst for a final 1X concentration of both reagents (1µg/mL) and placed in a 37°C/5% CO_2_ incubator for 1 hour following 72 hours of drug exposure. In spheroid culture, cells were treated with 5X (PI) for a final 1X concentration (1µg/mL) or 500x CytoToxic Green (Promega) for a final 1X concentration in the well. PI/CytoToxic green fluorescent images were acquired using the BioTek Cytation 5 (Agilent) with Gen 5 imaging software (Agilent) taken at 4x.

Cell counts were quantified using the Gen 5 imaging segmentation software. Cell death was quantified using two distinct CellProfiler pipelines (www.cellprofiler.org) for automated image analysis of monolayers or spheres. In short, the CellProfiler pipelines: 1) outlined the brightfield sphere or outlined all DAPI positive objects (monolayer); 2) calculated all objects with an RFP (propidium iodide) or GFP (CytoToxic Green) fluorescent intensity above a designated threshold for positivity; 3) saved values. The area of all fluorescent objects was then added together for total PI+ or GFP+ area in a given image and divided by total, cumulative nuclei area (Hoechst, monolayer) or total sphere area (brightfield, spheroid)

### Cell cycle analysis

In brief, FUCCI (mKO2-hCdt1 and mAG-hGeminin) expressing HCC827 monolayer cells were treated with afatinib (100nM, 20nM, 4nM, 0.8nM, 0.16nM, 0.032nM) 24 hours after plating. Following 72 hours of treatment, cells were analyzed by immunofluorescence microscopy using the BioTek Cytation 5 (Agilent) with Gen 5 imaging software (Agilent) taken at 4X. Post-imagining cellular analysis was performed using the Gen 5 data collection and analysis software (Agilent) by thresholding the RFP or GFP channels to meet the criteria for active G1 (Cdt1-RFP) or S/G2/M (Geminin-GFP).

### RT-qPCR analysis

Total RNA was isolated from cell lines using an RNeasy kit (Qiagen). Equal concentrations of total RNA were reverse transcribed using High-Capacity RNA-to-cDNA kit (ThermoFisher Scientific). The relative expression of *CTGF, CYR61*, and *TRAIL* was analyzed using a SYBR Green Real-Time PCR kit (Thermo) with an Applied Biosystems QuantStudio 6 Flex Real-Time PCR System thermocycler (Thermo). Variation was determined by the ΔΔCT method with GAPDH mRNA levels as an internal control. Data plotted as log_2_(ΔΔCT). The primers used were as follows:

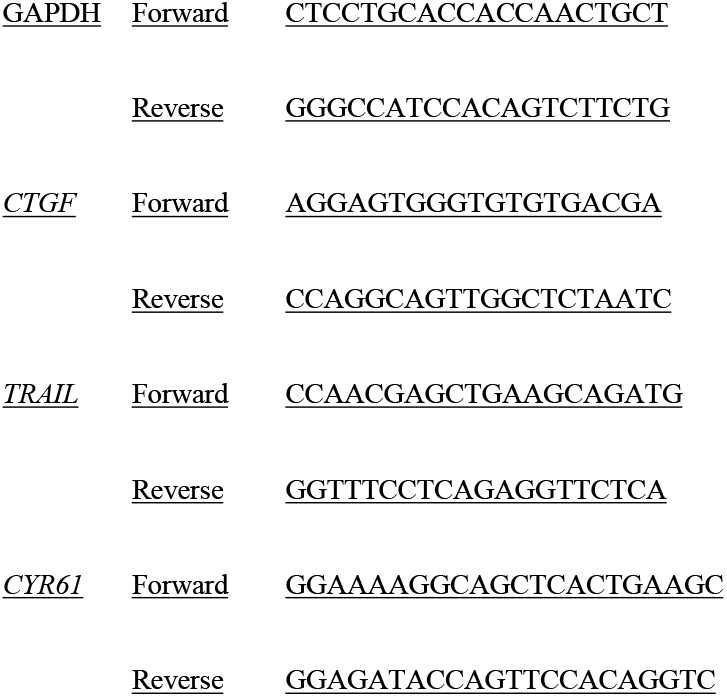

### Statistics

A *P* value less than .05 was considered significant, a *P* value greater than .05 was not significant (ns).

* *P* < 0.05, ** *P* < 0.01, *** *P* < 0.001, **** *P* < 0.0001

### Study approval

All murine experiments were approved by the Institutional Animal Care and Use Committee of UCSF.

### Data availability

Values for all data points in graphs are reported in the Supporting Data Values file.

## Methods Procedures References

### BioRender (Graphical Abstract): BioRender.com

#### Cell-profiler

Stirling, D.R., Swain-Bowden M.J., Lucas A.M., Carpenter, A.E., Cimini B.A^†^. and Goodman A^†^. (2021). CellProfler 4: improvements in speed, utility and usability. *BMC Bioinformatics* (22, 433). doi.org/10.1186/s12859-021-04344-9

#### StarDist Plugin

Schmidt, U., Weigert, M., Broaddus, C., & Myers, G. (2018). Cell Detection with Star-Convex Polygons. *Lecture Notes in Computer Science* (pp. 265–273). Springer International Publishing. doi:10.1007/978-3-030-00934-2_30

#### qPCR ΔΔCT

Livak, K.J. and Schmittgen, T.D. (2001). Analysis of Relative Gene Expression Data Using Real-Time Quantitative PCR and the 2^-ΔΔCT^ Method. *Methods* (pp. 402-408). Elsevier. https://doi.org/10.1006/meth.2001.1262

## Results

### EGFRi treatment of NSCLC xenografts selects for persister cells with predominantly nuclear YAP expression

Persister cells have been operationally defined as the surviving tumor cells after acute treatment with EGFR inhibitors (29). First, we sought to characterize the persister cells following acute EGFRi treatment compared to vehicle control. EGFR mutant NSCLC HCC827 (EGFRdel19) xenografts were generated in mice and treated short-term with afatinib, a second-generation, irreversible pan-HER kinase inhibitor (20mg/kg daily for 5 days), resulting in a rapid ~ 90% reduction in tumor volume compared to starting volume. During the same time period, vehicle-treated control tumors maintained growth and demonstrated increased volume (**Figure 1A**). Previous studies have implicated YAP as a potential mechanism of resistance to prolonged afatinib treatment or resistance (22,23), leading us to question if YAP was involved in resistance to even a short (5-day) treatment. Tumors harvested from afatinib- and vehicle-treated mice on day 5 of treatment were evaluated by immunofluorescence microscopy analysis of YAP localization in pan-cytokeratin positive cells (**Figure 1B**). Persistent cells within the remnant tumors of afatinib-treated mice consisted of ~60% pan-cytokeratin positive epithelial cells with nuclear YAP, compared to ~10% nuclear YAP of vehicle-treated tumors (**Figure 1C**). Afatinib-treated HCC827 persister cells also exhibited significantly decreased mitotic activity as indicated by phospho-Histone H3 staining (pHH3), consistent with prior work (29), but retained mutant EGFR as delineated by EGFRdel19 staining (**Supplemental Figure 1, A and B**). These data demonstrate an acute persistence phenotype in EGFRi-treated HCC827 xenografts characterized by YAP nuclear localization and activation driven by EGFRi.

**Figure 1.**
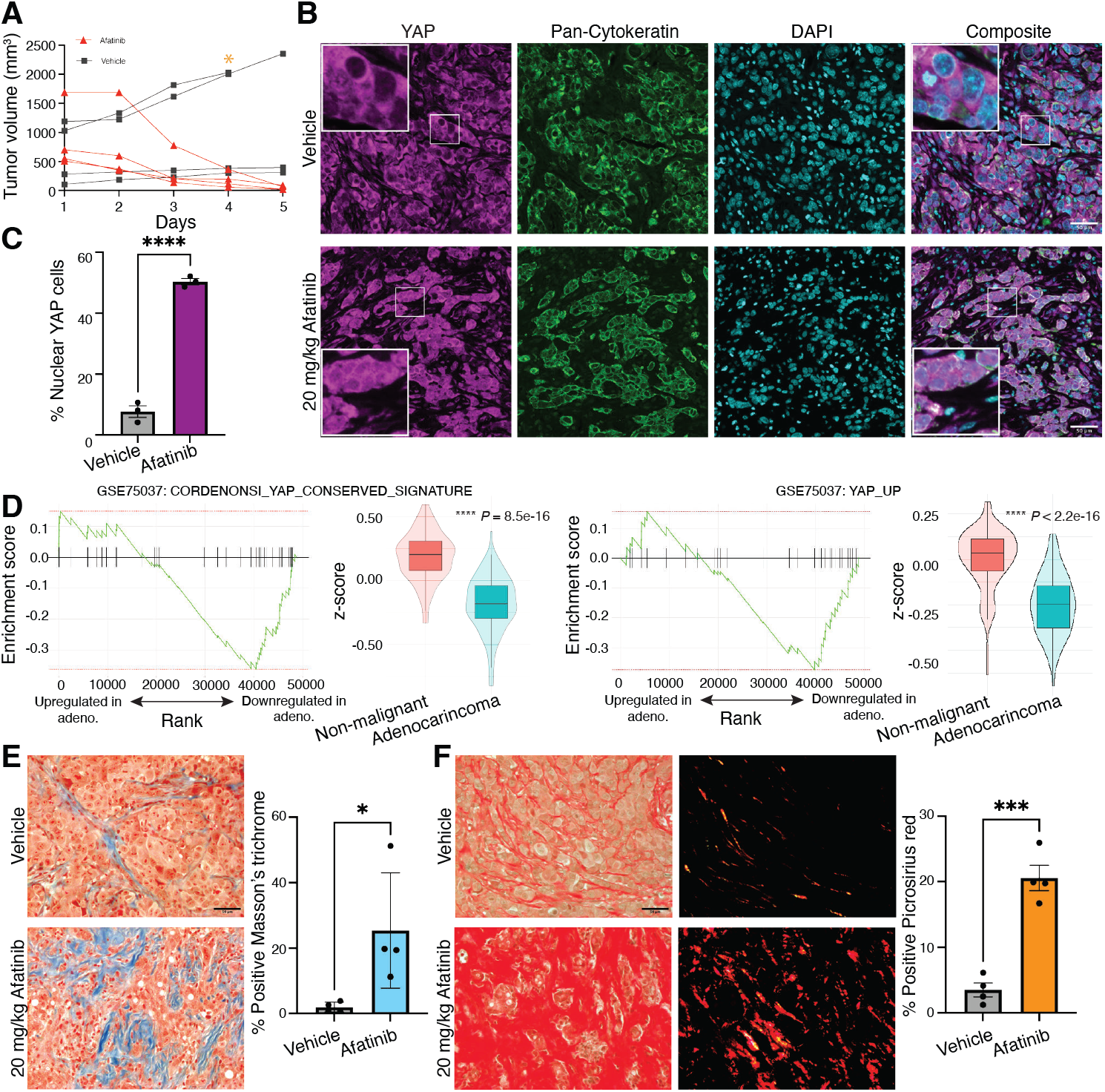
A persister phenotype in afatinib-treated tumor xenografts is driven by YAP activation. **(A)** Mice harboring HCC827 tumor xenografts were treated with 20mg/kg afatinib for 5d (*n* = 4) or vehicle (*n* = 4). **(B)** Immunofluorescence images of YAP, pan-cytokeratin, and DAPI in afatinib-treated remnant and vehicle tumors (magnification, 20x). **(C)** Quantification of nuclear YAP in vehicle- or afatinib-treated HCC827 xenografts. **(D)** Gene-set enrichment (GSEA) of YAP signatures from differential expression of 83 matched sets of patient adenocarcinoma or non-malignant tissue using YAP_conserved_signature [Normalized Enrichment Score = −1.76, p = 0.0039] (left) or YAP_up [Normalized Enrichment Score = −1.69, p = 0.0068] (right) gene signature in the ranked genes. Average Z-scores for each gene set were compared between conditions using a t-test [*P* = 8.5e-16 and *P <* 2.2e-16, respectively]. **(E)** Representative images of Masson’s trichrome and quantification of percentage area coverage of collagen for vehicle-treated tumors and afatinib-treated remnant tumors are shown (magnification, 10x). **(F)** Representative images of Picrosirius red staining and quantification of percentage area coverage per field of view (magnification, 10x). All scale bars, 50 μm. Data are shown as means ± s.e.m. (*n* = 3 biological replicates). * *P* < 0.05, *** *P* < 0.001, **** *P* < 0.0001 as determined by a two-tailed *t*-test.

Our findings in the xenograft studies indicate that, in the absence of treatment, YAP is predominantly localized to the cytoplasm in most cells, suggesting that this pathway is not active in the majority of tumor cells prior to treatment. We next queried if YAP activation is similarly downregulated in primary human cancers in the absence of treatment. We assessed the differential expression of YAP activity in 83 matched patient lung adenocarcinoma samples compared to adjacent non-malignant lung tissue (GSE75037)(37) and found that both the YAP conserved gene signature (38) and the YAP Up gene signature (39) were significantly downregulated in the adenocarcinoma compared to normal adjacent lung tissue (**Figure 1D**). These data suggest YAP is not intrinsically active in bulk lung tumors and indicate tumor cells that survive acute EGFRi therapy upregulate YAP. Given the mechanosensitive nature of YAP activity, we hypothesized that changes in cell-ECM dynamics following tumor reduction from acute EGFRi therapy could facilitate YAP nuclear sequestration after treatment. To investigate EGFRi-mediated changes to the ECM, we stained control and afatinib-treated tumor sections with Masson’s Trichome and Picrosirius red to evaluate total and fibrillar collagen ECM components following afatinib treatment (**Figure 1, E and F**). These data suggest that even short-term EGFRi treatment elicits an acute persistence phenotype in HCC827 xenografts characterized by YAP nuclear localization in DTPs embedded in dense, fibrotic collagen matrix.

### Spheroid culture sensitizes NSCLC cells to EGFRi-mediated apoptosis

Nuclear YAP was associated with dense ECM following EGFRi treatment, but not in vehicle treated HCC827 xenografts or transcriptional pathway activation in untreated NSCLC patient samples. We, therefore, hypothesized that the distinct stages of NSCLC tumor cells, from EGFRi-sensitive to DTPs, could be better modeled using distinct spheroid and monolayer culture contexts, respectively. We predicted that differences in culture conditions known to affect YAP activity—such as cell shape, substrate stiffness, and cell-cell adhesion (18,40,41)—in standard monolayer versus spheroid cell cultures would result in varying sensitivity to EGFRi based on culture context.

To test this, we generated spheroids from the HCC827 (EGFRdel19) cell line in ultra-low adhesion plates (lysates collected at 96 hr) or standard monolayer cell culture plates (lysates collected at 48 hr). First, we sought to characterize the distinct culture conditions. Baseline EGFR and EGFR phosphorylation (pEGFR) were significantly altered in HCC827 spheroids compared to monolayers, as monolayer cells displayed robust EGFR and pEGFR relative to total protein that was absent in spheroid culture (**Figure 2A**). Ponceau S staining intensity was utilized as a total protein loading control due to significant and unexpected differences in the protein levels of standard loading controls (42,43), including Vinculin and Histone H3, that were not reflective of total protein (**Supplemental Figure 2, A-D**). Actin and tubulin were excluded as potential loading controls due to distinct actin and tubulin organizational differences in monolayer compared to spheroid culture (44–46). Based on literature identifying a YAP-EGFR positive feedback loop in cervical cancer (47), we next queried whether monolayers and spheroids demonstrated differential YAP levels. While total YAP protein remained similar between monolayer and spheroids, phosphorylated YAP-S127 (pYAP) was significantly upregulated in spheroids compared to monolayer. Due to differential EGFR and pEGFR levels in monolayers compared to spheroids, we explored whether the different culture contexts would affect acute response to EGFRi. Surprisingly, dose-dependent relative ATP assays following 72 hr. treatment showed HCC827 spheres treated with afatinib demonstrated increased drug sensitivity relative to monolayer counterparts, as indicated by increased percent maximum ATP inhibition in spheroid versus monolayer culture (**Table 1, Figure 2B**). To ensure that spheroid cell culture is not inherently more sensitive to afatinib exposure and that afatinib selectively targets EGFRmut cells, we treated A549 (EGFR WT) spheroid and monolayer cell culture with afatinib. As expected, dose-dependent relative ATP assays confirm that A549 cells were not sensitive to afatinib-mediated inhibition when grown as spheroids (**Supplemental Figure 3A**). As an experimental control to confirm EGFR-mutated spheres are not inherently more sensitive to all chemotherapeutic compounds, HCC827 and A549 (EGFR WT) monolayer or sphere cultures were treated with cisplatin, a platinum based alkylating agent that targets actively dividing cells (48). HCC827 and A549 spheres exhibited increased resistance to cisplatin compared to monolayer cultures (**Figure 2C** and **Supplemental Figure 3B**), similar to prior reports (35). Thus, we conclude that the dose-dependent ATP inhibition observed in HCC827 spheres upon EGFRi is a specific response, rather than a general sensitivity produced by spheroid culture.

**Table 1.** Cell line EC50 and % Max Inh for targeted therapies.

| Cell line | Inhibitor | 72 hr Treatment |  |  |  |
| --- | --- | --- | --- | --- | --- |
|  |  | Monolayer (n=3) |  | Spheroid (n=3) |  |
|  |  | EC50<sup>A</sup> | % Max Inh.<sup>B</sup> | EC50 | % Max Inh. |
| HCC827 |  |  |  |  |  |
|  | Afatinib (nM) | 2.36 nM | 55.84 % | 1.52 nM | 89.94 % |
|  | Cisplatin (μM) | 4.55 μM | 88.98 % | --<sup>C</sup> | 0.00<sup>D</sup> |
| A549 |  |  |  |  |  |
|  | Afatinib (nM) | 10.11 nM | 30.00 % | 0.01 nM | 14.80 % |
|  | Cisplatin (μM) | 4.55 μM | 88.98 % | 75.33 μM | 86.13 %<sup>E</sup> |
| H-1975 |  |  |  |  |  |
|  | Osimertinib (nM) | 3.17 nM | 60.71% | 4.00 nM | 93.00 % |
| HCC60 |  |  |  |  |  |
|  | ARS-1620 (μM) | 7.7 μM | 87.13 %<sup>F</sup> | 1.62 μM | 89.49 % |
| H358 |  |  |  |  |  |
|  | ARS-1620 (μM) | 0.32 μM | 86.18 % | 0.23 μM | 79.94 % |
| RPMI-7951 |  |  |  |  |  |
|  | Vemurafenib (μM) | 9.46 μM<sup>G</sup> | 86.45 %<sup>H</sup> | 0.08 μM<sup>I</sup> | 79.97 % |
| HCC827-Control |  |  |  |  |  |
|  | Afatinib (nM) | N/C | N/C | 1.21 nM | 74. % |
| HCC827-YAP<sup>S127A</sup> |  |  |  |  |  |
|  | Afatinib (nM) | N/C | N/C | 1.27 nM | 61.79 % |
| H358-Control |  |  |  |  |  |
|  | ARS-1620 (μM) | N/C | N/C | 0.47 μM | 78.51 % |
| H358-YAP<sup>S127A</sup> |  |  |  |  |  |
|  | ARS-1620 (μM) | N/C | N/C | 1.18 μM | 85.24 % |

N/C, not calculated. <sup>A</sup>EC50 was calculated using GraphPad PRISM 9 nonlinear regression (curve fit) parameters. <sup>B</sup>% Max Inh. Was calculated using GraphPad PRISM 9 nonlinear regression fit parameters to determine the best-fit bottom value. The best-fit bottom value was then subtracted from 100 to yield % Max Inh. <sup>C</sup>Value was unstable. <sup>D</sup>Value was negative and rounded to 0.00. <sup>E</sup>Bottom best fit value was unstable, mean plotted value was used and manually calculated. <sup>F</sup>Bottom best fit value was negative, mean plotted value was used and manually calculated. <sup>G</sup>EC50 calculated from 5-fold dilution series of vemurafenib (.0064 μM – 20 μM). <sup>H</sup>Bottom best fit value was negative, mean plotted value was used and manually calculated. <sup>I</sup>EC50 calculated from 5-fold dilution series of vemurafenib (.01 μM – 20 μM).

**Figure 2.**
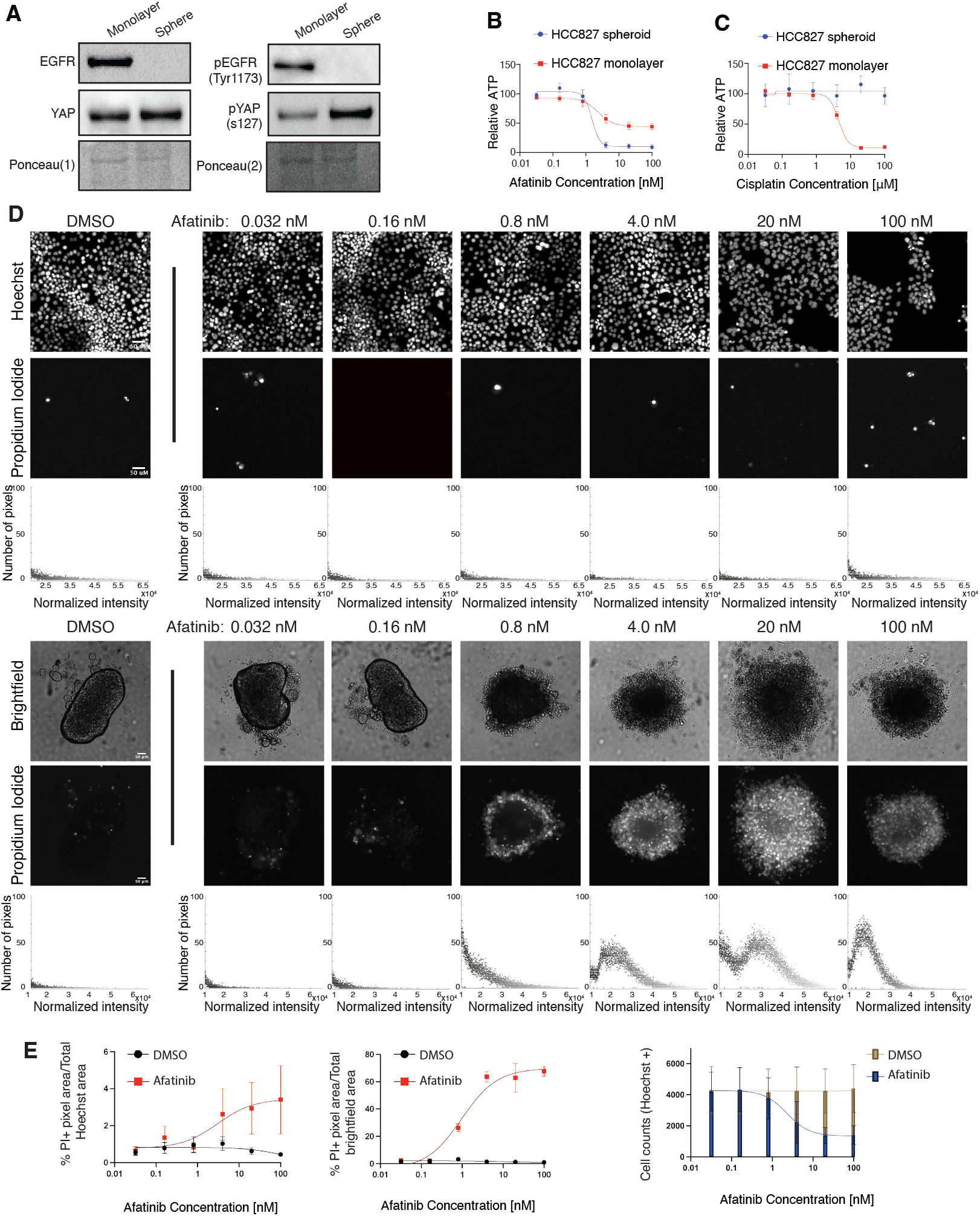
EGFR-mutated NSCLC spheroids, but not monolayers, acutely undergo massive apoptosis in response to EGFRi. **(A)** HCC827 spheroids (96 hr) and monolayers (48 hr) were harvested for western blot analysis against indicated proteins. Ponceau S was used as a total protein loading control. **(B)** Dose response curves for HCC827 cells cultured as spheroids or monolayers and treated with afatinib. Curves normalized to DMSO. LogEC50 (ns) *P* value = 0.4 between monolayer (EC50 = 2.36 nM) and spheroid (EC50 = 1.52 nM). * *P* value = 0.0199 between monolayer (Bottom value = 44.16 relative ATP) and spheroid (Bottom value = 10.06 relative ATP). **(C)** Dose-response curves for HCC827 cells cultured as spheroids or monolayers and treated with cisplatin. Curves normalized to 0.9% NaCl vehicle. **(D)** Representative Hoechst (monolayer) or brightfield (spheres) and corresponding Propidium Iodide fluorescence of DMSO- or afatinib-treated HCC827 monolayers and spheroids at indicated doses (magnification, 4x). All scale bars, 50 μm. Total, normalized RFP+ fluorescence pixels across each representative image shown. (E) Dose-response curves of % Propidium Iodide pixels / total Hoechst (monolayer) or **(F)** brightfield (spheroid) pixel area in DMSO- or afatinib-treated HCC827 monolayers and spheroids. (*n* = 3 biological replicates). **(G)** Quantification of total cell numbers across indicated DMSO- or afatinib-treated HCC827 monolayer cultures. Relative ATP was measured by a Cell Titer Glo assay. Data are shown as means ± s.e.m. (*n* = 3 biological replicates). * *P* < 0.05, ** *P* < 0.01, **** *P* < 0.0001 as determined by Extra sum-of-squares F Test; n.s., not significant.

**Figure 3.**
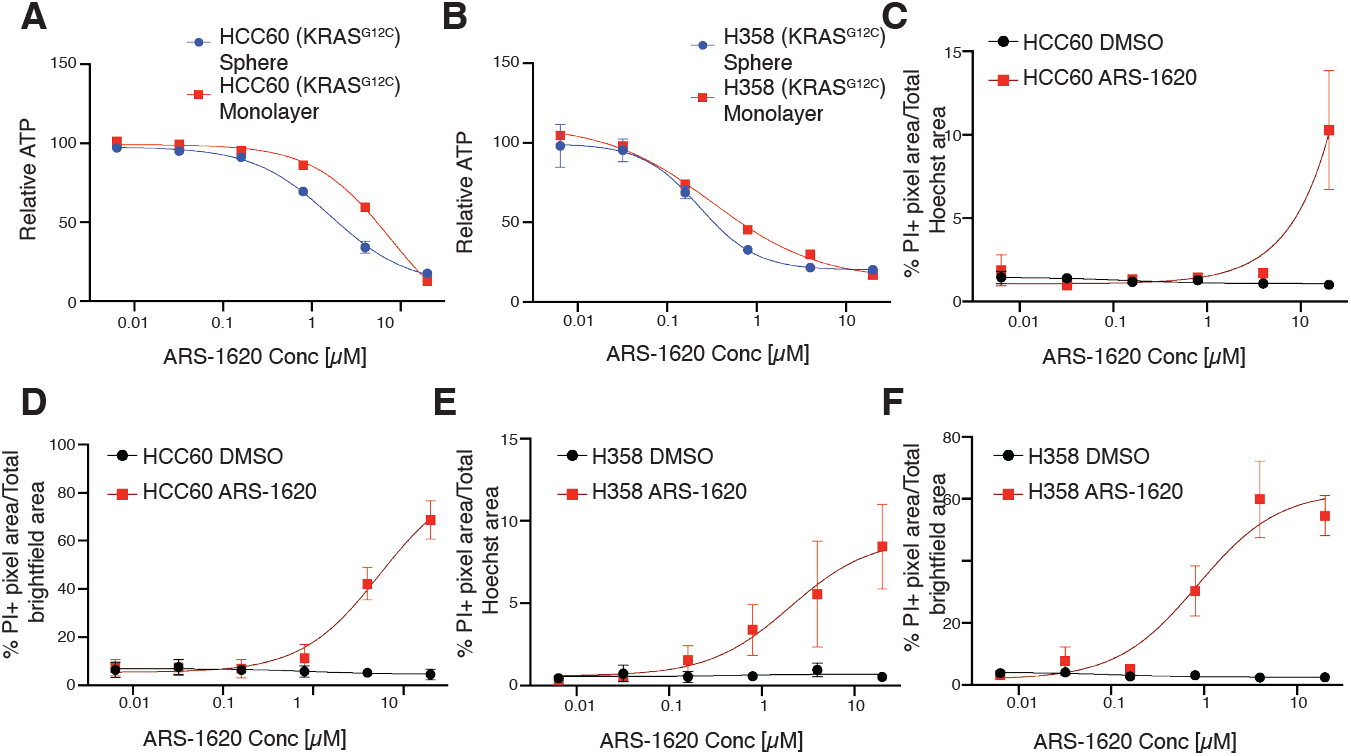
Distinct KRASG12C-mutant cell lines exhibit varying EC50 values in response to KRASG12C inhibition. **(A)** Dose response curves of HCC60 spheroids or monolayers treated with ARS-1620 for 72h. Relative ATP was measured by a Cell Titer Glo assay. Curves normalized to DMSO. LogEC50 **** *P* value = < 0.0001 between monolayer (EC50 = 7.70 µM) and spheroid (EC50 = 1.62 µM). **(B)** Dose response curves of H358 spheroids or monolayers treated with ARS-1620 for 72h. Relative ATP was measured by a Cell Titer Glo assay. Curves normalized to DMSO. LogEC50 (ns) *P* value = 0.415 between monolayer (EC50 = 0.32 µM) and spheroid (EC50 = 0.23 µM). **(C-D)** Dose-response curves of % Propidium Iodide pixels / total Hoechst (monolayer) or brightfield (spheroid) pixel area in DMSO- or ARS-1620-treated HCC60 monolayers and spheroids. **(E-F)** Dose-response curves of % Propidium Iodide pixels / total Hoechst (monolayer) or brightfield (spheroid) pixel area in DMSO- or ARS-1620-treated H358 monolayers and spheroids. Data are shown as means ± s.e.m. (*n* = 3 biological replicates). **** *P* < 0.0001 as determined by Extra sum-of-squares F Test; n.s., not significant.

Since relative ATP assays do not distinguish between cell death and or cell arrest, we next sought to determine whether afatinib elicited a cytostatic or cytotoxic response depending on culture conditions. We first treated monolayer (plated 24 hr) and sphere cultures (plated 96 hr) with afatinib or vehicle, then stained both monolayer or sphere cultures with propidium-iodide (PI) after 72 hr treatment. Relative to treated monolayer cultures, treated HCC827 spheres demonstrated massive cell death, as indicated by increasing PI uptake in spheres when treated with increasing doses of afatinib compared to DMSO. This is in stark contrast to monolayer, which demonstrates minor dose-dependent PI uptake compared to DMSO (**Figure 2, D and E**). Notably, EGFRi-treated HCC827 monolayers demonstrate ~50% decreased cell number relative to control despite minimal observed apoptosis at 72 hr relative to spheroid culture (**Figure 2F**), suggesting a predominant cytostatic response. To test if monolayer cell culture was primarily undergoing cell cycle arrest, we generated FUCCI (49) labeled HCC827 monolayers. We observed that HCC827 monolayers undergo robust G1 proliferative arrest in a dose-dependent manner after EGFRi treatment (**Supplemental Figure 4, A-C**). Thus, spheroid culture enforces a cell death phenotype while monolayer leads to tumor cell survival but G1 proliferation arrest. These data are reminiscent of our *in vivo* HCC827 xenograft models, where EGFRi-naïve tumor cells respond robustly to acute EGFRi treatment, while DTP cells embedded in the ECM show limited mitotic cells compared to vehicle control.

**Figure 4.**
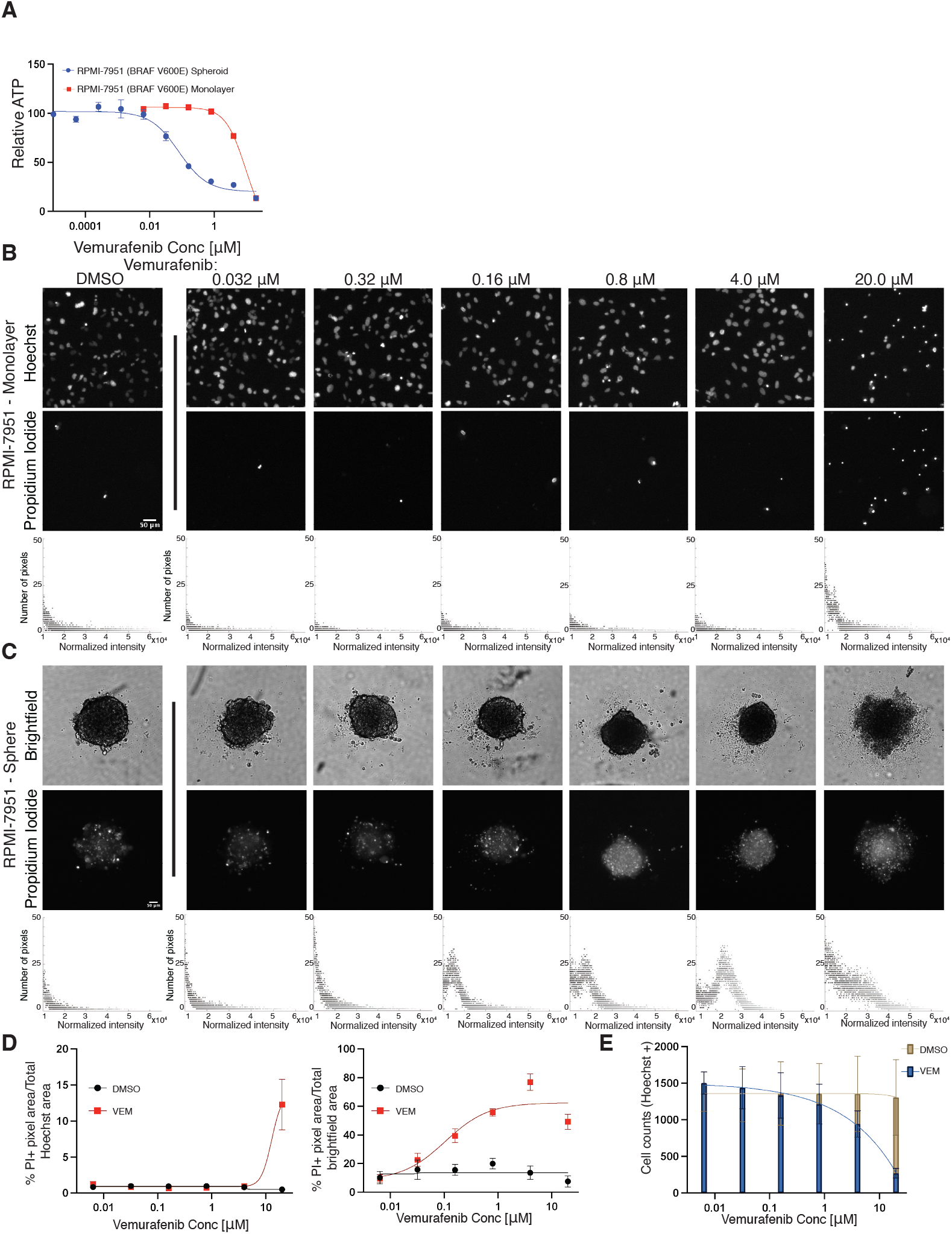
Vemurafenib resistant BRAF^V600E^ mutant line, RPMI-7951, is sensitive to vemurafenib treatment in spheroid culture. **(A)** Dose response curves of RPMI-7951 spheroids or monolayers treated with vemurafenib for 72h. Relative ATP was measured by a Cell Titer Glo assay. Curves normalized to DMSO. LogEC50 **** *P* value = < 0.0001 between monolayer (EC50 = 9.46 µM) and spheroid (EC50 = 0.08 µM). **(B-C)** Representative Hoechst (monolayer) or brightfield (spheres) and corresponding Propidium Iodide fluorescence of DMSO- or vemurafenib treated RPMI-7651 monolayers and spheroids at indicated doses (magnification, 4x). Total, normalized RFP+ fluorescence pixels across each image shown. **(D)** Dose-response curves of % Propidium Iodide pixels / total Hoechst (monolayer) or brightfield (spheroid) pixel area in DMSO- or vemurafenib-treated RPMI-7951 monolayers and spheroids. **(E)** Quantification of total cell numbers across indicated DMSO- or vemurafenib-treated RPMI-7951 monolayer cultures. Data are shown as means ± s.e.m. (*n* = 3 biological replicates). **** *P* < 0.0001 as determined by Extra sum-of-squares F Test.

### Enhanced sensitivity of EGFR-mutated NSCLC spheroids to EGFRi extends across multiple agents

We further investigated whether differential sensitivity is observed in NSCLC lines with other EGFR mutations by testing osimertinib, a third-generation, irreversible EGFRi that targets both EGFR activating mutations and inhibits the EGFR T790M resistance mutation (50). Similar to our HCC827 results, H-1975 spheres treated with osimertinib for 72 hr demonstrated increased drug sensitivity and massive cell death compared to monolayer counterparts (**Table 1, Supplemental Figure 5, A-C**). Additionally, H-1975 monolayers demonstrated ~70% reduction in cell numbers and minimal percent PI staining (**Supplemental Figure 5D**), confirming that osimertinib likely induces cytostatic effects in monolayer cultures and cytotoxic in spheroid cultures.

**Figure 5.**
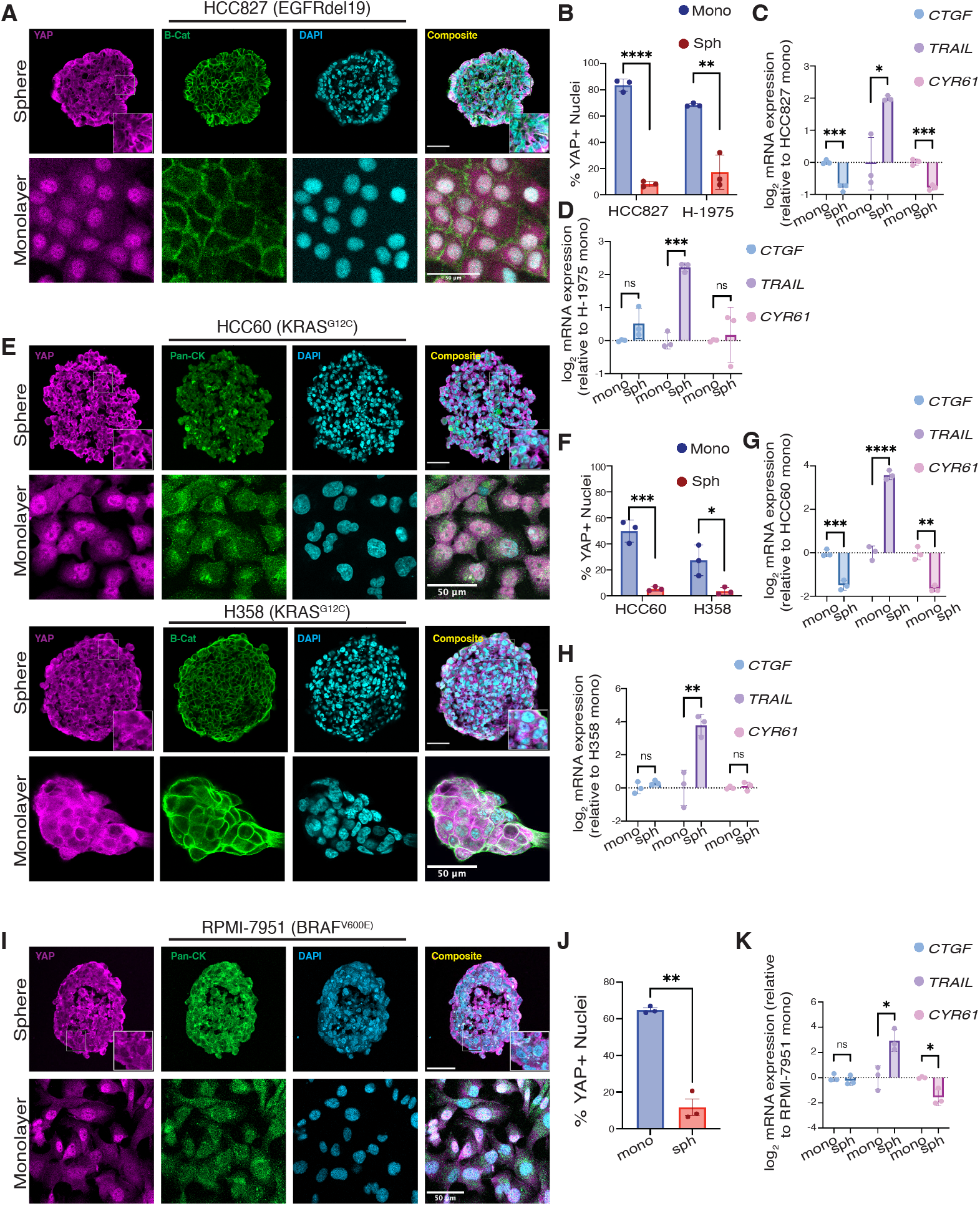
YAP nuclear translocation mediates resistance to EGFR/RAS/RAF pathway inhibitors. **(A)** Immunofluorescence images of YAP, β-catenin, and DAPI in EGFR mutant sphere (magnification, 20x) and monolayer (magnification, 10x) cultures. All scale bars 50μm. **(B)** Quantification of nuclear YAP to DAPI ratios in HCC827 and H1975 adherent monolayers and spheroids. Data are shown as means ± s.d. **(C-D)** qPCR analysis (*CTGF, TRAIL*,*CYR61*) in EGFR mutant spheroids normalized to monolayers. **(E)** Immunofluorescence images of YAP, pan-Cytokeratin (HCC60) or β-catenin (H358), and DAPI in KRAS^G12C^ mutant sphere and monolayer cultures (magnification, 20x). All scale bars 50μm. **(F)** Quantification of nuclear YAP to DAPI ratios in HCC60 and H358 adherent monolayers and spheroids. Data are shown as means ± s.d. **(G-H)** qPCR analysis (*CTGF, TRAIL, CYR61*) in KRAS mutant spheroids normalized to monolayers. **(I)** Immunofluorescence images of YAP, pan-cytokeratin, and DAPI in RPMI-7951 (BRAF^V600E^ mutant) sphere and monolayer cultures (magnification, 20x). All scale bars 50μm. **(J)** Quantification of nuclear YAP to DAPI ratios in RPMI-7951 adherent monolayers and spheroids. Data are shown as means ± s.d. **(K)** qPCR analysis (*CTGF, TRAIL, CYR61*) in BRAF mutant spheroids normalized to monolayers. Data are shown as means ± s.d. (*n* = 3 biological replicates). * *P* < 0.05, ** *P* < 0.01, *** *P* < 0.001, **** *P* < 0.0001 as determined by a two-tailed *t*-test; n.s., not significant.

### KRAS^G12C^ and BRAF^V600E^ mutant spheroids demonstrate increased sensitivity to KRAS G12Ci and BRAF^V600E^ inhibition compared to monolayer conditions

As both EGFR and KRAS are integral to the RAS-RAF-MEK-ERK signaling cascade, we next investigated whether inhibiting other downstream components of EGFR in this pathway would lead to increased inhibitor sensitivity in spheroid culture. We first selected two mutant KRAS^G12C^ cell lines, H358 and HCC60, and performed dose-dependent viability assays with G12Ci (ARS-1620). HCC60 spheres were more sensitive to ARS-1620 than monolayers (**Table 1, Figure 3A**), whereas H358 sphere and monolayer cultures were similarly sensitive to ARS-1620 (**Table 1, Figure 3B**). Likewise, quantitative analysis of monolayers and spheres stained with PI emphasized sensitivity differences between HCC60 and H358-while HCC60 monolayers only showed robust cell death at the maximum concentration of ARS-1620 used (**Figure 3C**), the HCC60 spheroids maintained a sigmoidal curve for %PI positivity, indicating cells were dying at lower concentrations of drug compared to monolayer (**Figure 3D**). In contrast, both H358 spheroids and monolayer culture demonstrate dose-dependent cell death when exposed to ARS-1620, as indicated by the sigmoidal curves of %PI+ positivity (**Figure 3, E and F**). These data provide evidence of a culture context-specific resistance mechanism protecting HCC60 monolayer culture from G12Ci-mediated cell death that is not present in HCC60 spheres or the H358 cell line, regardless of H358 culture context. Our results agree with published findings of KRAS^G12C^ mutant lines based on differential sensitivity to ARS-1620 in monolayer vs sphere culture, which has reported similar EC50 values for the H358 cell line in response to ARS-1620 treatment in monolayer and spheres (9).

To extend our analysis beyond the primary NSCLC EGFR and KRAS driver mutations, we additionally looked at monolayer and sphere dose response differences in a cell line carrying a BRAF^V600E^ mutation, which is present in less than 5% of NSCLC patients (1). The sole immortalized lung adenocarcinoma cell line carrying a BRAF^V600E^ mutation found in the ATCC catalog, HCC364, does not form spheres in cell culture (data not shown). We therefore tested a melanoma cancer cell line, RPMI-7951 (BRAF^V600E^), as a representative tumor cell line for the BRAF^V600E^ mutation. We then tested if vemurafenib, a selective inhibitor used in the clinic to treat patients with BRAF^V600E^ mutant kinase melanoma (51), demonstrated differential efficacy in RPMI-7951 spheroids compared to monolayer cell culture. RPMI-7951 cells in monolayer culture were resistant to vemurafenib at concentrations above 3 µm, in agreement with previously reported findings (52,53) (**Table 1, Figure 4A**). We observed, however, that RPMI-7951 spheres lack this resistance, as demonstrated by the robust cell death observed after 72h of treatment at much lower concentrations in the absence of collagen scaffolding (53) (**Table 1, Figure 4, B and C**). Quantifying RFP+ positive pixels, indicative of PI uptake, in RPMI-7951 spheroids compared to monolayer culture following 72 hr vemurafenib treatment confirms the increased sensitivity of spheroid cell culture. Spheroids displayed robust, dose-dependent cell death, whereas monolayer culture only demonstrated increased apoptosis and reduced cell counts at the highest dose tested. (**Figure 4, B-E**). Intriguingly, RPMI-7951 resistance to vemurafenib has been reported to be caused by a MAP3K8 amplification in RPMI-7951 cells (52). We confirm resistance in monolayer culture, similar to other mutants in the EGFR/RAS pathway; however, spheroid culture unmasks increased sensitivity.

Taken together, these data demonstrate that short-term (acute) responses of EGFRi-, G12Ci-, and V600Ei-targeted therapies in NSCLC and melanoma can be specific to culture context: in several of the cases examined, cells grown in spheres showed increased sensitivity to drug in comparison to cells grown in monolayer. Based on clinical results emphasizing the robust, though transient, response of EGFRi and G12Ci in patient cohorts, our data suggest that spheroid cell culture better mimics initial patient response to therapy than standard monolayer culture, while monolayer models DTP cells.

### Monolayer culture promotes YAP nuclear localization, mediating resistance to EGFRi-induced cell death in NSCLC

Given our in vivo HCC827 xenograft findings highlighting differential YAP localization in vehicle vs. afatinib-treated tumors, we hypothesized YAP may be cytoplasmic in spheroids in which integrin-extracellular matrix interactions are disengaged, predisposing spheroids to EGFRi, G12Ci, and V600Ei-mediated apoptosis. Since mechanical and morphological cues can modulate YAP localization and activity (18), we assayed YAP localization in EGFR, G12C, and V600E-mutated spheroids and monolayers. Strikingly, YAP localized to the nucleus in both EGFR mutated cell lines as monolayers, but predominantly localized to the cytoplasm in spheroids (Figure 5A and Supplemental Figure 6A). These data indicate that YAP nuclear localization corresponds to the acute persistence phenotype in EGFRi-treated monolayers (**Figure 2** and **Supplemental Figure 5**), reminiscent of the persister cells identified in the HCC827 xenograft model (**Figure 1**).

**Figure 6.**
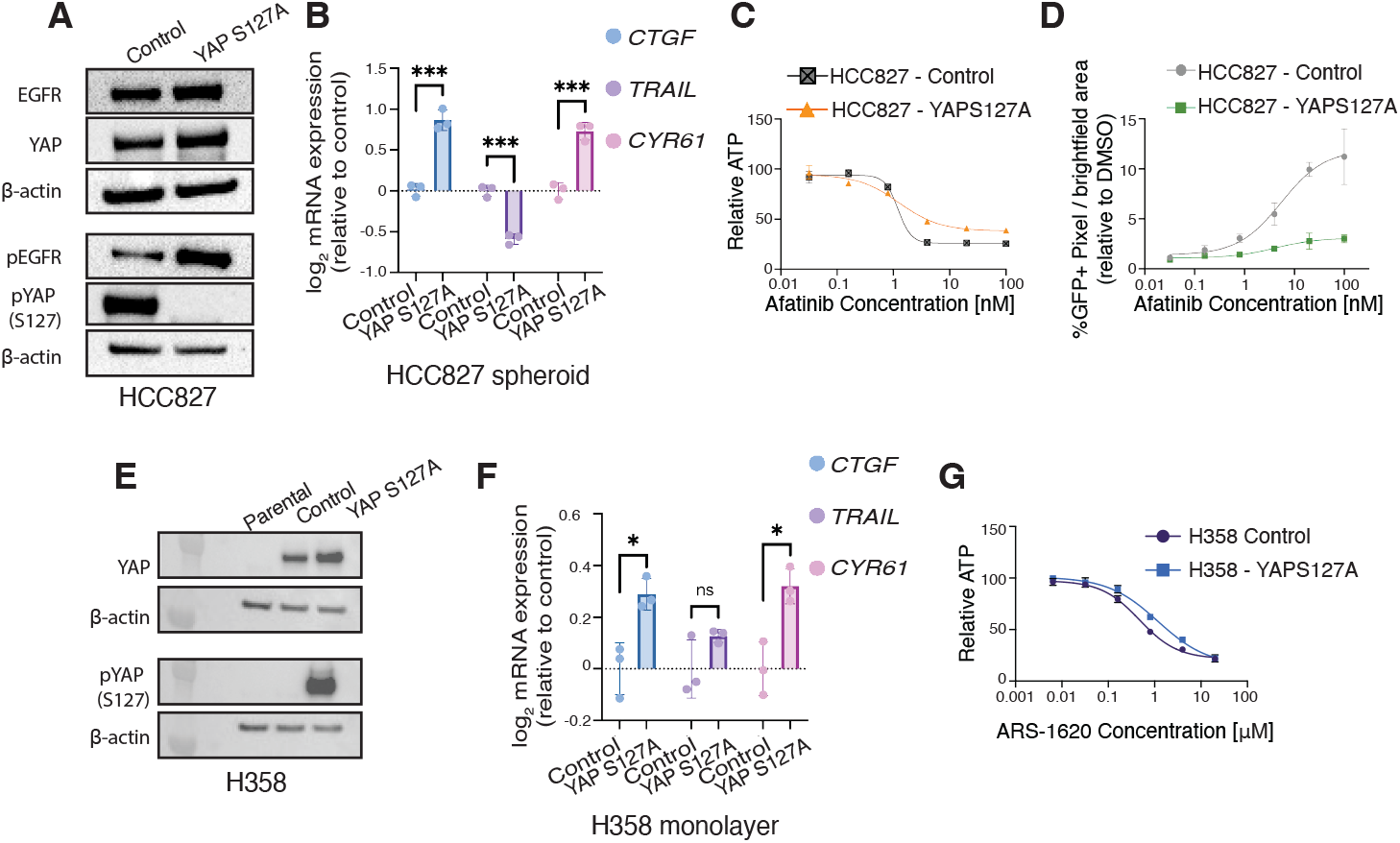
Increased YAP activity is sufficient to protect HCC827 spheroids from afatinib-mediated cell death. **(A)** HCC827-control and HCC827-YAP^S127A^ monolayers (48 hr) were harvested for western blot analysis against indicated proteins. **(B)** qPCR analysis (*CTGF, TRAIL, CYR61*) in HCC827-YAP^S127A^ spheroids normalized to HCC827-control. Data are shown as means ± s.e.m. (*n* = 3 biological replicates). **(C)** Dose response curves for HCC827-control compared to HCC827-YAP^S127A^ cells cultured as spheroids and treated with afatinib for 72 hr. Relative ATP was measured by a Cell Titer Glo assay. Curves normalized to DMSO. LogEC50 (ns) *P* value = 0.91 between control (EC50 = 1.21 nM) and YAP^S127A^ (EC50 = 1.27 nM). * *P* value = 0.021 between HCC827-control (Bottom value = 25.98 relative ATP) and HCC827-YAP^S127A^ (Bottom value = 38.21 relative ATP). **(D)** Dose-response curves of % GFP pixels / total brightfield pixel area in HCC827-control and HCC827-YAP^S127A^ normalized to DMSO % GFP pixels / total brightfield pixel area. Data are shown as means ± s.e.m. (*n* = 3 biological replicates). **(E)** H358-parental, H358-control, and H358-YAP^S127A^ monolayers (48 hr) were harvested for western blot analysis against indicated proteins. **(F)** qPCR analysis (*CTGF, TRAIL, CYR61*) in H358-YAP^S127A^ monolayers normalized to H358-control. Data are shown as means ± s.e.m. (*n* = 3 biological replicates). **(G)** Dose response curves for H358-control compared to H358-YAP^S127A^ cells cultured as spheroids and treated with ARS-1620 for 72 hr. Relative ATP was measured by a Cell Titer Glo assay. Curves normalized to DMSO. LogEC50 ** *P* value = 0.005 between H358-control (EC50 = 0.47 µM) and YAP^S127A^ (EC50 = 1.18 µM). * *P* < 0.05, ** *P* < 0.01 as determined by Extra sum-of-squares F Test; n.s., not significant.

Based on cytoplasmic sequestration of YAP across our spheroid models, we next examined if YAP transcriptional activity was altered in NSCLC spheroids compared to respective monolayers. We queried the expression of three established YAP transcriptional targets (upregulation of *CTGF* and *CYR61*; downregulation of *TRAIL* (37,39,54)) and found the expression of these genes to be altered as expected for HCC827 spheres compared to monolayer (**Figure 5C**), consistent with increased YAP activity in monolayers. Interestingly, for H-1975, *CTGF*, and *CYR61* gene expression were not significantly altered between monolayer and spheres, an unexpected result, given that YAP is predominantly nuclear in monolayer by immunofluorescence analysis (**Figure 5D**). Since *CTGF* and *CYR61* were identified as primary YAP targets in monolayer cultures, these data question if YAP is a universal transcriptional activator of *CTGF* and *CYR61* in spheroid culture across all cancer cell types.

After identifying YAP localization differences in the EGFRmut cell lines between sphere and monolayer culture, we hypothesized that the KRAS^G12C^ cell lines would not be uniform in their YAP localization. As expected, H358 cells, which did not demonstrate robust difference in sensitivity between monolayer and spheroid culture to ARS-1620 (Figure 3), demonstrated only a modest increased YAP nuclear localization in monolayer compared to spheroid. In contrast, HCC60 cells, which exhibit a robust delta between the EC50 of ARS-1620 of spheroid and monolayer culture, displayed YAP predominantly localized to the nucleus in monolayer culture and sequestered to the cytoplasm in spheroid culture (**Figure 5, E and F**). Differential gene expression of *CTGF, CYR61*, and *TRAIL* were as expected for HCC60, with significant downregulation of *CTGF* and *CYR61* and upregulation of *TRAIL* in spheroid culture compared to monolayer, consistent with YAP activity in monolayer culture (**Figure 5G**). Surprisingly, though *CTGF* and *CYR61* were not significantly altered in H358 spheroids relative to monolayer, *TRAIL* was significantly upregulated (Figure 5H).

Finally, we assessed differential YAP localization in the RPMI-7951 cell line between intrinsically vemurafenib-resistant monolayer culture and vemurafenib-sensitive spheroid culture. Consistent with previous reports implicating YAP activation in vemurafenib resistance (55), we observed monolayer cells had a significantly higher percentage of cells with nuclear YAP than spheroid culture. In contrast, spheroid culture demonstrated YAP predominantly sequestered to the cytoplasm (**Figure 5, I and J**). Despite the expected changes in *CYR61* and *TRAIL* expression between RPMI-7951 spheres and monolayer cultures, *CTGF* expression remained unchanged, contrary to what might be expected given YAP’s nuclear localization in monolayer cultures (**Figure 5K**). Interestingly, in all five lines tested, *TRAIL* expression was significantly upregulated in spheroid culture compared to monolayer. Future studies would be needed to elucidate if *TRAIL* is universally upregulated in spheroid compared to monolayer culture independent of YAP activity, as all cell lines queried for this work demonstrated robust *TRAIL* upregulation, regardless of *CTGF* or *CYR61* expression.

Taken together, these data suggest that culture context may non-genetically alter localization in a cell-line specific manner. In our studies, YAP nuclear localization correlates to relative drug efficacy and whether a drug exhibits cytostatic or cytotoxic effects. These results highlight the need for cell culture systems beyond standard monolayer culture, as consequential alterations of YAP activity depending on the culture method may unexpectedly confound studies assessing drug sensitivity and resistance.

### Upregulation of YAP activity confers increased resistance to EGFRi and KRAS^G12C^i-targeted therapies

Given our data demonstrating that differential YAP localization between spheroid and monolayer cultures correlates to altered drug sensitivity, we hypothesized that promoting YAP nuclear localization could render cells resistant to EGFRi even in spheroid culture. To test this hypothesis, we generated HCC827 cells that overexpress YAP^S127A^ or YAP^WT^ (YAP overexpressing control). The YAP^S127A^ mutant increases YAP nuclear translocation by altering a canonical LATS1/2 phosphorylation site that targets YAP for proteasomal degradation in the cytoplasm, increasing available YAP protein for translocation to the nucleus (56–58). Though baseline YAP protein levels remained equal between YAP^S127A^ and or YAP^WT^, HCC927-YAP^S127A^ cells had dramatically less YAP S127 phosphorylation compared to HCC827-control cells, as expected (**Figure 6A**). Baseline EGFR was also equivalent between control and HCC827-YAP^S127A^, though there was a notable increase in pEGFR in the HCC827-YAP^S127A^ cells. These results were surprising since previous work has demonstrated that YAP overexpression and YAP^S27A^ mutants express pEGFR compared to parental cells at similar levels (59), yet our data suggests a positive feedback loop between YAP activity and pEGFR. We next examined YAP activity for HCC827-YAP^S127A^ cells and HCC827-control cells grown in spheroid culture. qPCR analysis validated the increased activation of YAP in the HCC827-YAP^S127A^ cells relative to HCC827-control, with increased expression of *CTGF* and *CYR61* and the downregulation of *TRAIL* (**Figure 6B**). Sensitivity to afatinib as measured by dose-response assays of HCC827-YAP^S127A^ spheres treated with Afatinib compared to HCC827-control spheres confirmed our hypothesis, demonstrating that HCC827-YAP^S127A^ spheres had reduced % maximum inhibition compared to controls and protected spheres from robust cell death relative to control spheres (**Table 1, Figure 6, C** and **D and Supplemental Figure 7A**). These results indicate that increased nuclear YAP promotes resistance in EGFR mutated NSCLC to afatinib in spheroid culture, highlighting how YAP activity can confer resistance even in previously sensitive HCC827 spheroid culture.

After determining that increased YAP activity was sufficient to partially rescue HCC827-YAP^S127A^ spheroids from EGFRi-mediated cell death, we next investigated whether increased YAP activity could similarly protect H358 monolayer cultures from G12Ci-mediated cell death. H358 cells exhibited similar EC50 values between monolayer and spheroid cultures (**Figure 3B**), and qPCR analysis of *CTGF* and *CYR61* revealed comparable YAP activity levels in monolayers and spheroids (**Figure 5H**). As expected, both H358-control and H358-YAP^S127A^ cells displayed elevated basal YAP protein expression compared to H358 parental cells. H358-control cells, however, displayed robust phosphorylation of YAP at the S127 site, validating the YAP^S127A^ mutation (**Figure 6E**). qPCR analysis further confirmed increased YAP activity in the H358-YAP^S127A^ cells, with significantly higher expression of *CTGF* and *CYR61* in the H358-YAP^S127A^ compared to H538-control cells. Surprisingly, *TRAIL* gene expression remained unchanged in the H358-YAP^S127A^ mutant relative to H358-control cells, highlighting the need for further investigation into whether YAP activity is antagonistic to TRAIL expression only under specific cellular contexts. Dose-dependent assays confirmed that YAP activity protected H358-YAP^S127A^ cells from ARS-1620-mediated cell death, as evidenced by a significantly higher EC50 in H358-YAP^S127A^ cells compared to H358-control, indicating reduced sensitivity to the KRAS^G12C^ inhibitor (**Table 1, Figure 6G**). These results, taken together with the HCC827-YAP^S127A^ data, underscore the validity of our model (**Graphical Abstract**), in which induced YAP activity-whether through culture context for plasmid expression-can confer resistance to cells that were previously sensitive to these inhibitors. This suggests that increased YAP activity may play a critical role in modulating cellular responses to targeted therapies, potentially influencing the development of DTPs and permanent genetic resistance mechanisms in cancer cells.

## Discussion

Here, we have shown that acute driver oncogene inhibition in EGFR, KRAS, and BRAF-mutated NSCLC and melanoma have differential impacts dependent on culture condition. Our data suggest these discrepant responses are dictated by YAP localization in a cell-line specific manner. Cells cultured in standard monolayer versus spheroid culture alters YAP localization and activity by a non-genetic mechanism across multiple driver oncogenes. These findings are underscored by the YAP nuclear localization observed in remnant tumors following acute EGFRi-treatment for 5 days. Persister cells in our model are characterized by activation of YAP and are surrounded by a remodeled collagen-rich ECM. These studies underscore the necessity for considering diverse cell culture models beyond conventional monolayers to mitigate unforeseen YAP-related impacts on drug sensitivity and resistance studies and reveal a non-genetic mechanism of in vivo and in vitro resistance. Furthermore, this work reveals a non-genetic in vivo and in vitro resistance mechanism to diverse EGFR/RAS/RAF inhibitors following only acute treatment. Ultimately, these data provide a compelling model for studying the evolution of mutant NSCLC resistance to targeted inhibitors, in which spheroid cell culture more closely replicates treatment-naïve tumors, while monolayer culture inherently mimics DTPs.

The use of matrix-disengaged 3D spheroids in cell culture has become popular for modeling tumor biology due to its ability to maintain cell-cell contacts and preserve a multicellular architecture resembling patient tumors, offering a more physiologically relevant model than traditional monolayer cultures (33–36). This approach has proven particularly valuable in therapeutic development, enabling investigation into differential drug sensitivities between monolayers and spheroids (9,60). Indeed, our study on NSCLC showed decreased viability and increased cell death in monolayers compared to spheroids following cisplatin treatment, highlighting spheroid resistance to cytotoxic chemotherapy relative to monolayers (35). Inherent differences between spheroid and monolayer culture contexts also call into question whether canonical pathways are translatable between monolayer and sphere culture. In this work, for instance, canonical YAP targets *CTGF* (CCN2) and *Cyr61* (CCN1) correlated to YAP protein localization and subsequent activity by gene expression analysis in two cell lines tested, HCC827 and HCC60. *CTGF* and *Cyr61* gene expression levels, however, were unchanged in H-1975, H358, and RPMI-7951 (*Cyr61* was downregulated, as expected, for this line) spheres relative monolayers, even though YAP protein levels showed significant differences in nuclear localization between monolayer and spheres. Canonical YAP targets such as *CTGF* and *Cyr61* were identified in monolayer cultures, but variables in alternative culture conditions could alter the regulation of these targets. For example, hypoxic gradients intrinsic to spheroid culture (61) could alter these targets in spheroids, as *CTGF* expression has been shown previously to be repressed under hypoxic conditions in tubular epithelial cells (62). Future work will be needed to elucidate if canonical YAP targets, and downstream targets of other mechanosensitive transcription factors (MSTFs) (63), are universal between monolayer and spheroid culture.

Similarly, the observed discrepancies in basal EGFR and pEGFR protein levels between HCC827 monolayer and spheroid cultures raise important questions regarding the relationship between pEGFR protein expression and pathway dependency. Previous research in head and neck squamous cell carcinoma (HNSCC) has demonstrated that, while high levels of pEGFR generally correlate with cell line sensitivity to erlotinib, specific cell lines with low basal pEGFR expression exhibited comparable sensitivity (64). These findings align with our model, wherein spheroids display high dependence on EGFR autophosphoylation, as indicated by robust EGFRi-mediated cell death, despite lower pEGFR levels compared to their monolayer counterparts. This suggests that low basal pEGFR levels may not accurately reflect minimal pathway activation in these cells, and that pEGFR expression should not be considered a reliable biomarker for sensitivity to EGFRi.

Previous work identified YAP as an effector of resistance to apoptosis in EGFR-mutated NSCLC (22,23), BRAF-activating mutant melanoma (55,65), pancreatic adenocarcinoma driven by KRAS (66), and HER2-overexpressing breast cancer (67). Interestingly, forced YAP nuclear translocation using a non-phosphorylatable YAP^S127A^ mutant reduced HCC827 spheroid sensitivity to afatinib compared to YAP^wt^ overexpression. These data suggest that increased YAP activity protects from afatinib-mediated cell death, indicating that canonical YAP signaling can override morphological cues driving YAP localization in the context of NSCLC spheroids. These data support our findings that monolayer cell culture is more resistant to EGFRi, G12Ci, and BRAFi through non-genetic YAP activation inherently driven by culture context. Though activating mutations, such as YAP^S127A^, are rare in solid tumors (68,69), these results demonstrate that YAP activity can profoundly influence drug response, emphasizing the importance of understanding YAP activity as it relates to cell culture models and systems.

Our findings support previous reports on DTPs following EGFRi in patient samples. From those studies, DTP cells undergo G1 arrest following EGFRi to resist treatment-mediated cell death (29). Similarly, our studies investigating acute afatinib treatment in vivo resulted in near complete tumor regression, yet residual tumor cells that retained nuclear YAP were embedded in dense, highly cross-linked collagen and were predominantly cell cycle arrest. From these data and our results indicating mutant EGFR NSCLC cells undergo G1 cell cycle arrest upon acute afatinib treatment, we would speculate that the highly cross-linked collagen mimics ex vivo monolayer culture, in which cells are grown on stiff polystyrene plastic, driving nuclear YAP localization and causing non-genetic resistance to afatinib-mediated cell death.

Additionally, the data presented in this work suggest that combinatorial therapy of EGFRi and YAP inhibitors may provide synergistic benefits to eliminate persister cells following acute EGFRi treatment. Lee *et al*. reported that verteporfin and fluvastatin, two clinically available YAP/TAZ inhibitors, can resensitize gefitinib and afatinib-resistant H-1975 and HCC827 monolayer cultures, respectively, to EGFRi (23). Further work is necessary to determine the optimal timing for administering combinatorial YAP inhibition, as our results suggest that YAP inhibition may have limited efficacy when administered during acute EGFRi treatment when YAP activity is low.

In summary, we show that acute, 5-day in vivo treatment of NSCLC xenografts with EGFRi allows us to detect persister cells with nuclear YAP embedded in remodeled and dense collagen matrix. By comparing spheroid and monolayer cultures of common mutations identified in NSCLC, we have delineated YAP activation as a non-genetic resistance mechanism against EGFRi, G12Ci, and V600Ei-mediated apoptosis.

## Author Contributions

R.N. designed and conducted the experiments and wrote the manuscript. A.B. significantly contributed to study design, hypothesis generation, and intellectual input. S.D. assisted in qPCR studies and HCC827 and H-1975 relative ATP dose-response assays. M.K.H. performed Masson’s trichrome and picrosirius red staining, quantified Masson’s trichrome and picrosirius red staining in xenograft tissue, and revised the manuscript. V.S. performed differential YAP gene signature expression analysis of GSE75037 data. N.P.M. developed the YAP-S127A plasmid, provided valuable discussion, and contributed to manuscript revision. H.W. generated a YAP localization pipeline using the StarDist extension in QuPATH. H.G. supervised differential YAP gene signature analysis and provided valuable input. V.M.W. supervised Masson’s trichrome and picrosirius red staining and microscopy of xenograft tissue and provided scholarly input. D.V. assisted and supervised experimental design, provided intellectual input, and assisted in manuscript revision. A.G. supervised all studies and provided valuable discussion and scholarly input.

## Acknowledgements

This work was supported, in part, by the US NIH F31 CA265248 Predoctoral Individual National Research Service Award (R.N.), National Science Foundation Graduate Research Fellowships Program (R.N.), Molecular Pathology of Cancer training grant T32 CA177555-02 (A.B.), the Uniting Against Lung Cancer/Lung Cancer Research Foundation Legacy Program (A.B.), and a California Institute of Regenerative Medicine (CIRM) fellowship (A.B.), the Gazarian Endowment and Bechtle Family (A.G), The Mark Foundation (A.G) and (NIH R01-CA136717, R01-CA170447, 2R01EB028148 A.G.) and the Atwater Foundation (A.G.). We thank Dr. Erica Hutchins for her microscopy expertise and assistance.

---

**Figure S1.**
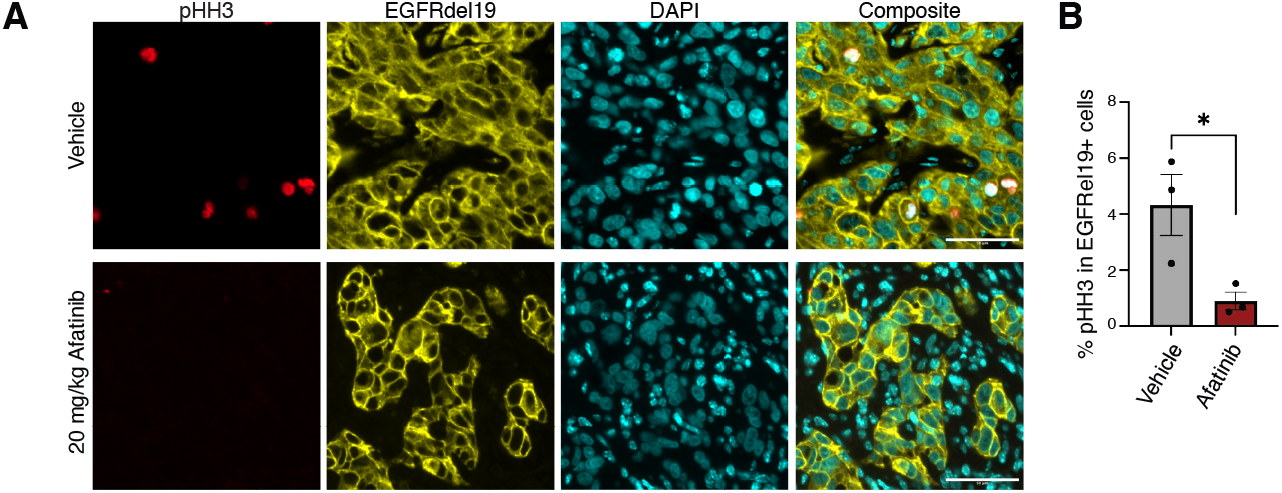
Afatinib-treated remnant tumors demonstrate reduced mitotic activity compared to vehicle-treated tumors. **(A)** Representative immunofluorescence images of vehicle or afatinib-treated remnant tumors stained with phospho-HistoneH3 (pHH3), EGFRdel19, or DAPI (magnification, 20x). **(B)** Quantification of pHH3 staining in the vehicle compared to afatinib-treated remnant tumors. All scale bars, 50 μm. Data are shown as means ± s.e.m. (*n* = 3 biological replicates). * *P* < 0.05 as determined by a two-tailed *t*-test.

**Figure S2.**
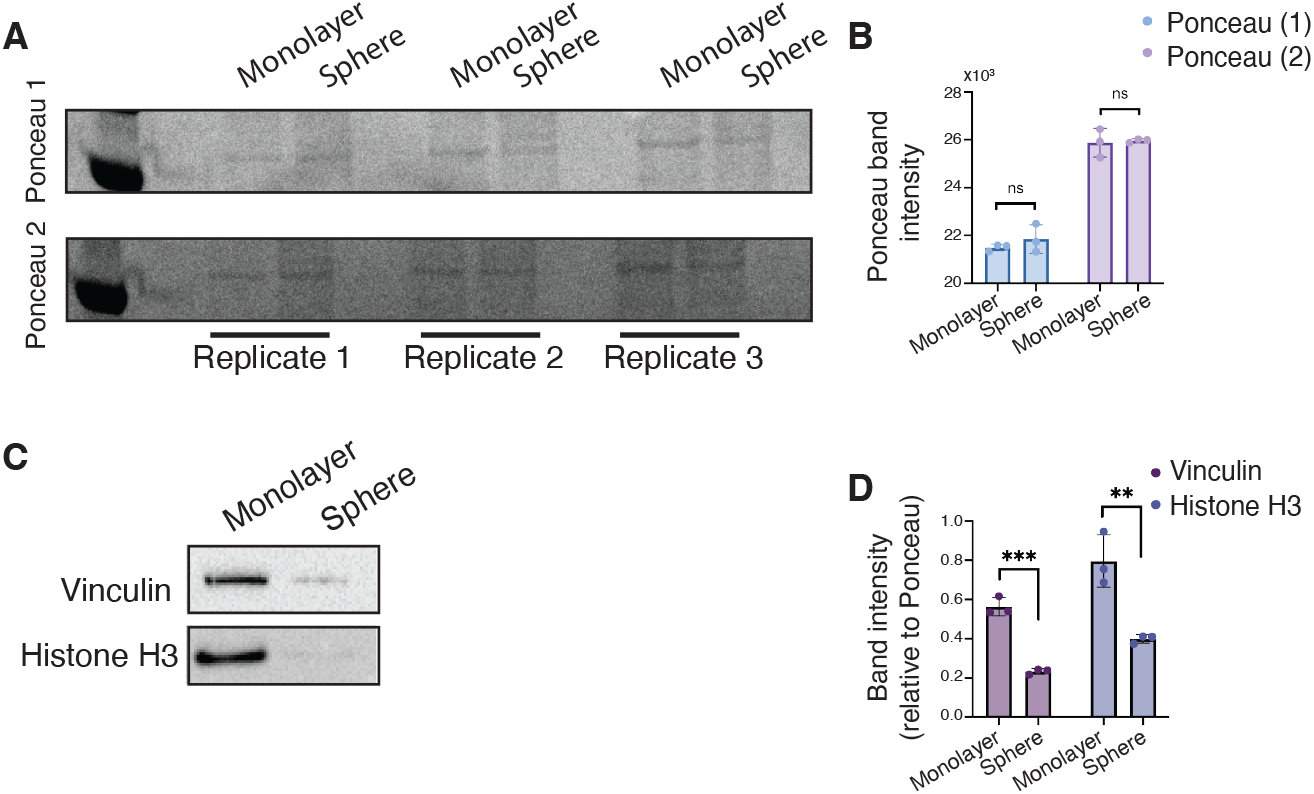
Standard loading controls for western blot analysis cannot be used for comparative analysis of monolayers and spheroids. **(A)** Ponceau staining of *n*=3 replicates of paired monolayer and spheroid protein lysate. **(B)** Quantification of ponceau band intensity. **(C)** Representative images of Vinculin and Histone H3 western blot bands in monolayer and sphere lysate. EGFR and pEGFR blots in Figure 2 were stripped and re-probed with vinculin. **(D)** Quantification of Vinculin and Histone H3 bands relative to Ponceau band intensity.

**Figure S3.**
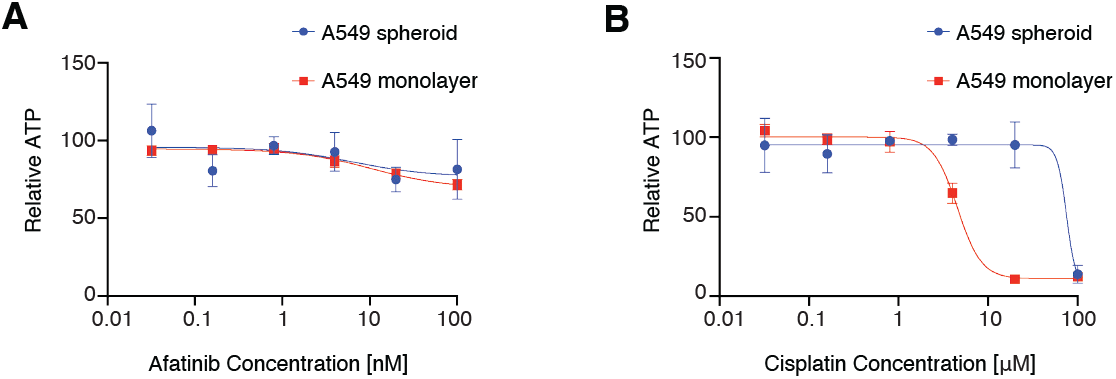
Cells do not demonstrate compromised overall fitness resulting from spheroid culture. **(A)** Dose-response curves for A549 cells cultured as spheroids or monolayers and treated with afatinib. Curves normalized to DMSO. LogEC50 \*\**P* value = 0.0023 between monolayer (EC50 = 4.55 μM) and spheroid (EC50 = 75.33 μM). **(B)** Dose-response curves for A549 cells cultured as spheroids or monolayers and treated with cisplatin. Curves normalized to 0.9% NaCl vehicle. Relative ATP was measured by a Cell Titer Glo assay. Data are shown as means ± s.e.m. (*n* = 3 biological replicates). * *P* < 0.05, ** *P* < 0.01, **** *P* < 0.0001 as determined by Extra sum-of-squares F Test; n.s., not significant.

**Figure S4.**
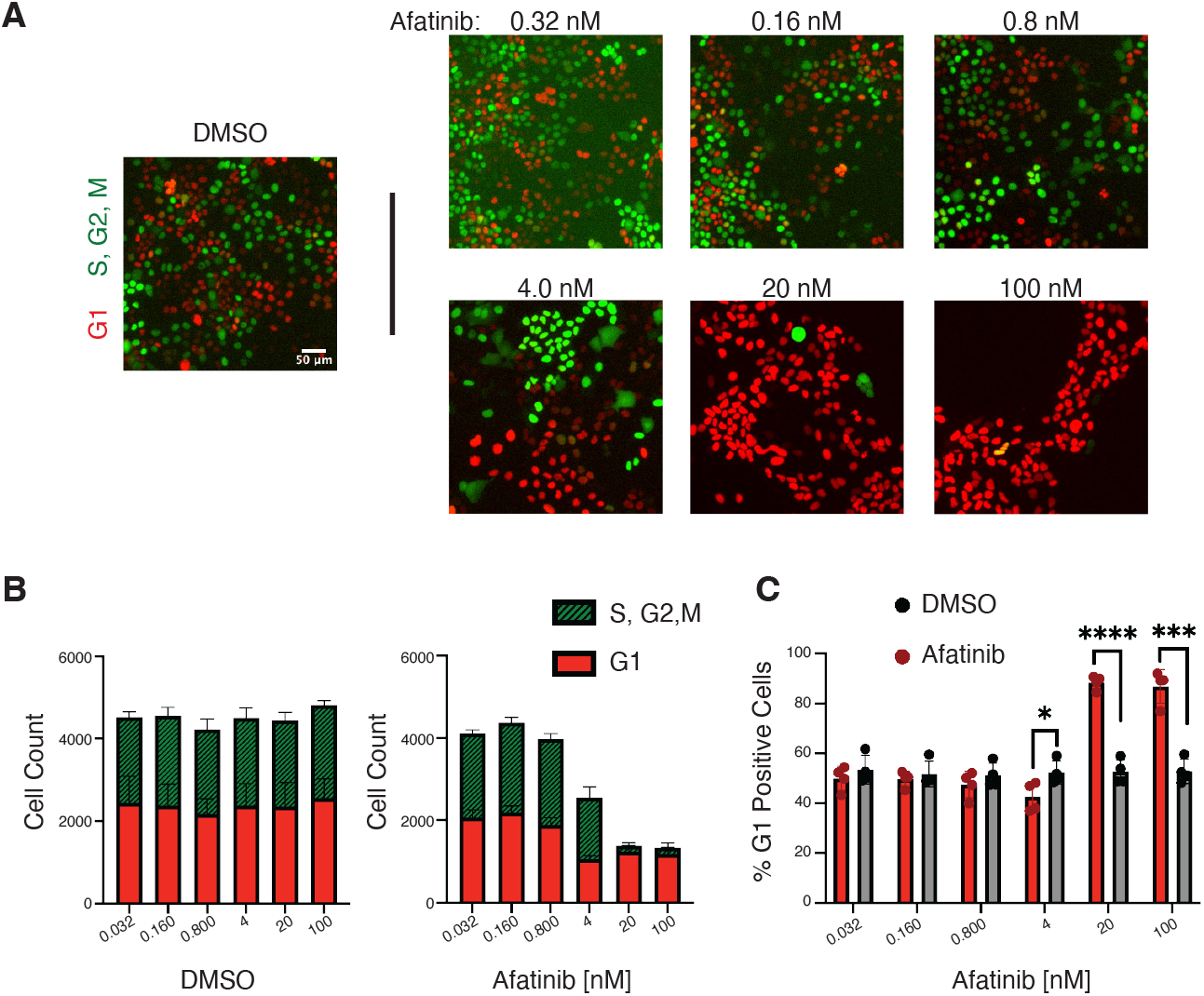
EGFR-mutated NSCLC monolayers undergo G1 arrest as an acute response to EGFR inhibition. **(A)** FUCCI (mKO2-hCdt1 and mAG-hGeminin) expressing HCC827 monolayers were treated with DMSO or afatinib at indicated doses for 72h and analyzed by immunofluorescence. Representative images shown (magnification, 4x). Scale bar 50μm. **(B)** Cell counts of mKO2-hCdt1 and mAG-hGeminin HCC827 expressing to quantify dose-dependent cell-cycle distribution from FUCCI expressing HCC827 monolayers following 72h of DMSO or afatinib treatment. **(C)** Percentage of mKO2-hCdt61 expressing cells (G1 cell cycle marker) over total cells. Data are shown as means ± s.e.m. (*n* = 4 biological replicates). * *P* < 0.05, *** *P* < 0.001, **** *P* < 0.0001 as determined by a two-tailed *t*-test.

**Figure S5.**
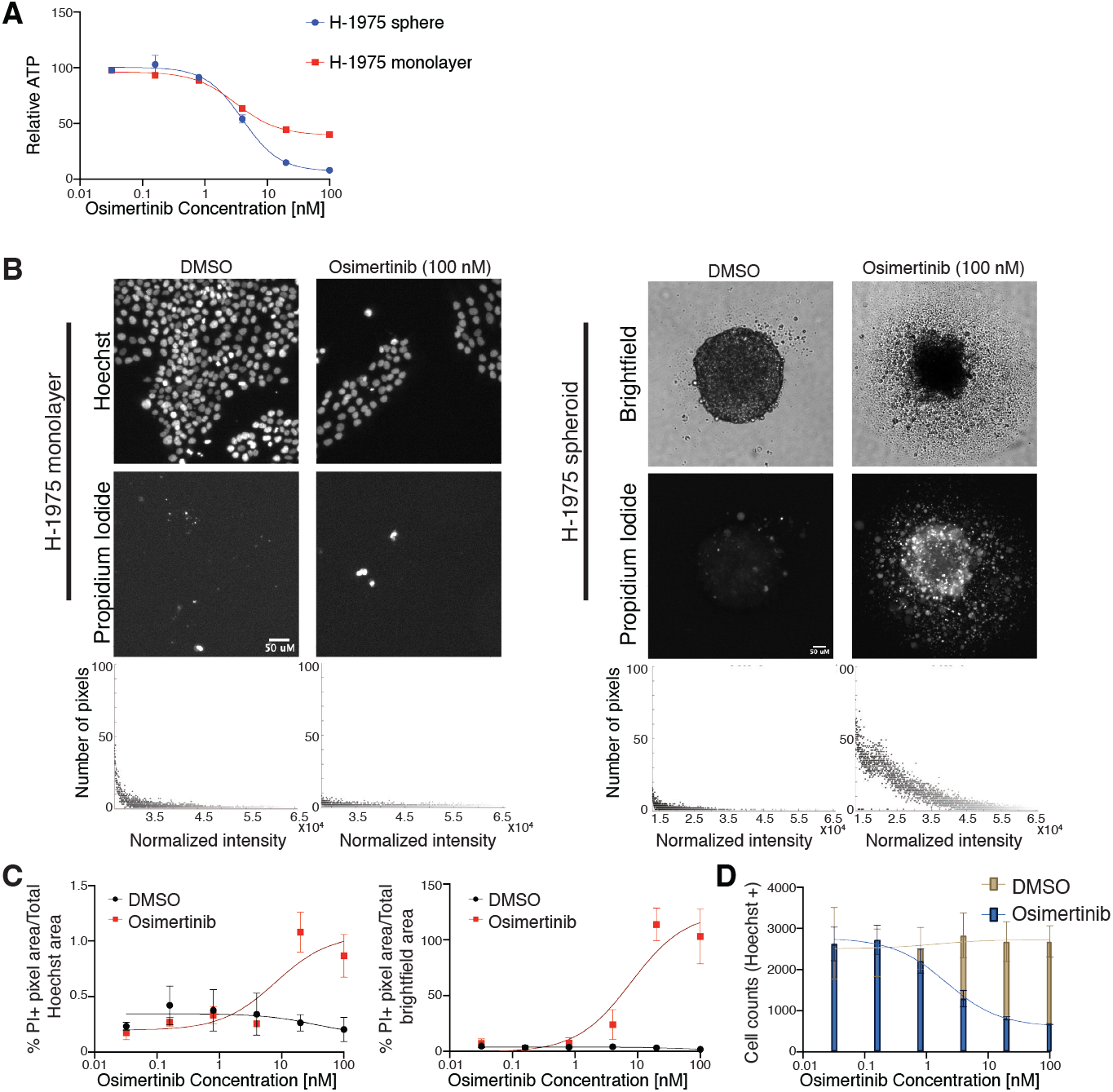
Differential sensitivity between EGFR-mutated spheroids and monolayer to EGFRi is not limited to a single agent or cell line. **(A)** Dose response curves of H-1975 spheroids or monolayers treated with osimertinib for 72h. Curves normalized to DMSO. Relative ATP was measured by a Cell Titer Glo assay. LogEC50 (ns) *P* value = 0.50 between monolayer (EC50 = 3.17 nM) and spheroid (EC50 = 4.00 nM). * *P* value = 0.023 between monolayer (Bottom value = 39.29 relative ATP) and spheroid (Bottom value = 7.01 relative ATP). **(B)** Representative Hoechst (monolayer) or brightfield (spheres) and corresponding Propidium Iodide fluorescence of DMSO- or 100 nM osimertinib-treated H-1975 monolayers and spheroids at indicated dosages (magnification, 4x). Total, normalized RFP+ fluorescence pixels across each image shown. **(C)** Dose-response curves of % Propidium Iodide pixels / total Hoechst (monolayer) or brightfield (spheroid) pixel area in DMSO- or osimertinib-treated H-1975 monolayers and spheroids. (*n* = 3 biological replicates). **(D)** Quantification of total cell numbers across indicated DMSO- or afatinib-treated HCC827 monolayer cultures. Relative ATP was measured by a Cell Titer Glo assay. Data are shown as means ± s.e.m. (*n* = 3 biological replicates). ns = not significant, * *P* < 0.05 as determined by Extra sum-of-squares F Test.

**Figure S6.**
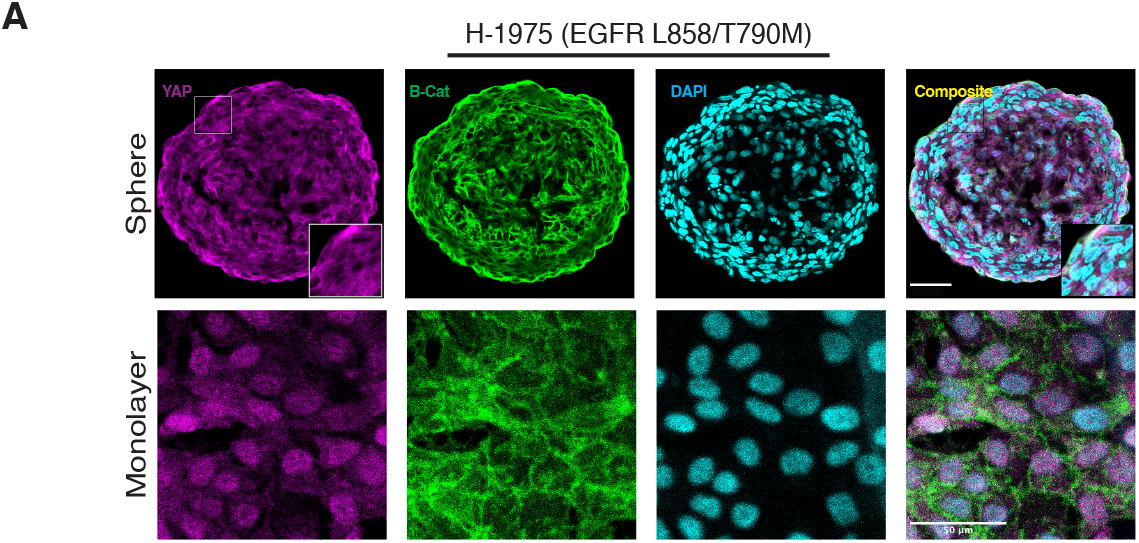
YAP exhibits cytoplasmic localization in H-1975 spheroids. **(A)** Immunofluorescence images of YAP, β-catenin, and DAPI in EGFR mutant sphere (magnification, 20x) and monolayer (magnification, 10x) cultures. All scale bars 50μm.

**Figure S7.**
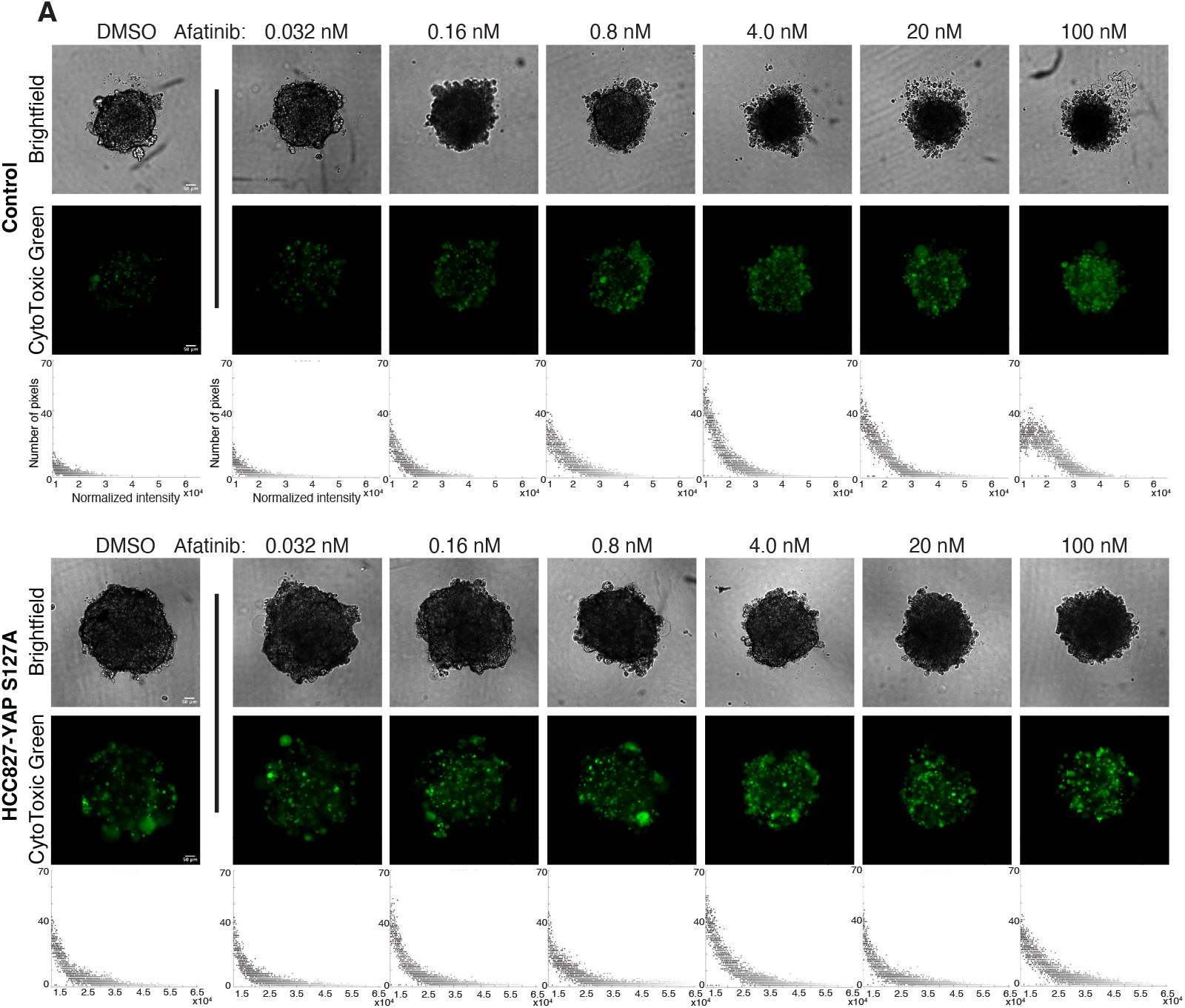
Dose-response assays using CytoToxic Green show HCC827-YAP^S127A^ cells are partially protected from EGFRi-mediated cell death. **(A)** Representative brightfield images and corresponding CytoToxic Green fluorescence of DMSO- or afatinib-treated HCC827-control and HCC827-YAP^S127A^ spheroids at indicated doses (magnification, 4x). Total, normalized green fluorescence pixels across each image shown.

